# Rare variants alter mitochondrial lipid homeostasis and neuronal excitability in PD patient-derived dopaminergic neurons

**DOI:** 10.64898/2026.04.10.717646

**Authors:** Federica Carrillo, Giorgio Fortunato, Arianna Coppola, Marco Ghirimoldi, Nwife Getrude Okechukwu, Vittoria Federica Borrini, Shahzaib Khoso, Antonietta Di Lorenzo, Mariarca Marciano, Giuseppe Giurin, Francesca D’amato, Maria Roberta Iazzetta, Cristina D’Aniello, Alessandro Fiorenzano, Teresa Nutile, Danilo Licastro, Sara Pietracupa, Nicola Modugno, Katiuscia Martinello, Sergio Fucile, Marcello Manfredi, Annalisa Fico, Teresa Esposito

## Abstract

Parkinson’s disease (PD) exhibits substantial genetic heterogeneity, yet how combinations of rare variants converge on disease-relevant cellular mechanisms remains unclear. Here, we generated human induced pluripotent stem cell-derived dopaminergic neurons from PD patients carrying rare variants in recently implicated genes and performed integrated electrophysiological, proteomic, lipidomic, and genetic analyses. Patient-derived neurons showed reduced membrane capacitance and altered action potential firing, indicating impaired intrinsic excitability and synaptic dysfunction, with marked variability across genetic backgrounds. Multi-omics profiling revealed dysregulation of mitochondrial function, glycolysis, and oxidative phosphorylation, accompanied by extensive lipid remodeling, including increased fatty acids, acylcarnitines, and sphingolipids, and reduced gangliosides. These alterations were more pronounced in neurons harboring specific variant combinations in KIF21B, SLC6A3, HMOX2, TMEM175, and AIMP2. Integrative analyses uncovered coordinated protein–lipid changes linking mitochondrial dysfunction and membrane homeostasis. Notably, Calpastatin and CXCR4 were consistently dysregulated across PD neurons. Genetic association analyses in independent cohorts identified PD-associated variants in genes encoding dysregulated proteins, supporting the functional relevance of these pathways. Overall, our results define convergent and variant-specific mechanisms underlying PD and highlight candidate biomarkers and therapeutic targets.

## Background

Parkinson’s disease (PD) is a neurodegenerative movement disorder caused by the inexorable and progressive loss of dopaminergic (DA) neurons in the Substantia Nigra pars compacta (SNpc), which accumulate intracellular α-synuclein oligomers [1]. PD is characterized by a slow progression with accumulation of various motor and non-motor disability in the affected individuals. Motor manifestations appear only after more than 50% of nigrostriatal dopaminergic neurons have degenerated, whereas non-motor features such as hyposmia, constipation, and sleep disturbances may arise during a prodromal phase (up to 15-20 years before motor onset), when underlying neurodegeneration is ongoing but clinical signs are not yet evident [2]. Current symptomatic therapies compensate the movement deficits efficiently only for some years before causing disabling dyskinesia [3]. Moreover, these therapies are unable to delay or halt the progressive neurodegenerative process. Parkinson’s disease is an age-related disease, with incidence and prevalence increasing steadily with age. The etiological factors underlying PD onset remain under investigation, with genetic factors estimated to account for approximately 25 % of the overall risk [1,4,5]. With the advancement of genome-wide association studies in large cohort of PD patients and controls, over 20 monogenic forms of PD have been described, and over 100 loci have been identified as risk factors for PD [4,6–8].

Monogenic variants with high or partial penetrance have been identified as causative of PD, in several genes including *SNCA*, *PARK2*, *PARK7*, *LRRK2*, *VPS35*, and *PINK1*, which collectively account for 5-10 % of all PD cases [9]. However, it is known that the spectrum of genetic variants underlying PD aetiology could range from rare variants with a considerable effect (fully or highly penetrant mutations in single genes) to genetic variants exerting only modest effects, which are relatively common in the general population. Furthermore, rare variants in Mendelian PD genes can also act as risk factors for late-onset, sporadic PD [10]. In this complex contest, a polygenic rare variant load model of inheritance has been identified in PD patients [11–13]. This model proposes that the co-inheritance of multiple rare variants in Mendelian genes may increase the risk of PD in a non-Mendelian fashion and influence both disease onset and clinical presentation [12,13].

A large body of studies have greatly advanced our understanding of the pathophysiological mechanisms underlying PD using cellular and animal models [14]. However, the marked clinical and genetic heterogeneity of PD limits the ability of any single experimental model to fully recapitulate the complexity of the human disease. In this context, patient-derived induced pluripotent stem cells (iPSCs) represent a powerful platform for investigating disease-relevant cellular phenotypes within defined genetic backgrounds. Notably, the differentiation of hiPSCs into midbrain dopaminergic neurons provides a particularly relevant system for modeling PD-associated alterations, including mitochondrial dysfunction, oxidative stress, impaired proteostasis, and synaptic alterations [15–17]. To date, iPSCs have been generated from patients with idiopathic PD (without a clear genetic cause) as well as from individuals carrying rare, highly penetrant mutations in dominant or recessive PD genes such as *SNCA, LRRK2, PARK7, PINK1*, and *GBA* [18–21]. Despite the substantial number of iPSC lines generated over the past decade, cellular models representing newly identified combinations of PD-associated genetic variants remain largely unavailable, limiting the investigation of their potential synergistic or modifying effects on disease pathogenesis [18–27].

Furthermore, the functional consequences of many of these rare variants remain poorly characterized, particularly within disease-relevant human cellular contexts. To address this gap, we generated a panel of patient-derived iPSC lines carrying highly penetrant combinations of genetic variants across eleven PD-associated genes: *AIMP2*, *ANKK1*, *HMOX2*, *KIF21B*, *LRRK2*, *RHOT2*, *SLC6A3*, *TMEM175*, *TOMM22*, *TVP23A* and *ZSCAN21*. These genes are implicated in key cellular pathways relevant to PD pathogenesis, including mitochondrial metabolism, oxidative stress, vesicular trafficking, microtubule dynamics, and lysosome-autophagy function [28–40].

We performed an extensive characterization of these newly generated iPSC lines, including analyses of pluripotency, chromosomal integrity, and differentiation potential. Our primary objective was to derive high-quality ventral midbrain dopaminergic progenitors (FOXA2+/LMX1+) and differentiate them into mature dopaminergic neurons. These neurons were subsequently subjected to functional electrophysiological assessments and integrated multi-omics analyses, including proteomics and lipidomics, to investigate the additive or synergistic effects of multiple PD-associated variants on neuronal function.

Our results disclosed pronounced functional alterations and different proteomic and lipidomic landscapes of patient-derived dopaminergic neurons harbouring distinct combinations of PD-related genetic variants. Finally, the integration of different omics layers led to the identification of novel modifiers factors for PD.

## Materials and Methods

### Study participants

PD patients used in this study were part of the PD biobank of IRCCS Neuromed/IGB-CNR. All the subjects were of European ancestry and were evaluated by qualified neurologists of the Parkinson Centre of the IRCCS INM Neuromed with a thorough protocol comprising neurological examination and evaluation of non-motor domains. Information about family history, demographic characteristics, anamnesis, and pharmacological therapy was also collected (the treatment of the PD groups consisted for the most part of a combination of levodopa and dopamine agonists) [12,13]. The Movement Disorder Society revised version of the Unified Parkinson’s Disease Rating Scale Part III (33 items, maximum score 132; hereafter called UPDRS) [41] and Hoehn e Yahr (HY) scales were used to assess clinical motor symptoms. These included language, facial expressions, tremor, rigidity, agility in movements, stability, gait and bradykinesia. Cognitive abilities were tested through an Italian validated version of the Montreal Cognitive Assessment (MoCA) [42]. Cognitive domains assessed include short-term memory, visuospatial abilities, executive functioning, attention, concentration, and working memory, language and orientation to time and place.

Non-motor symptoms were assessed through an Italian validated version of Non-Motor Symptoms Scale (NMS) for Parkinson Disease [43]. This scale tests 9 items, including cardiovascular domain, sleep/fatigue, mood/cognition, perceptual problems/hallucinations, attention/memory, gastrointestinal, urinary, sexual function, and ability to taste or smell.

Whole exome sequencing (WES) data were used to identify mutations/variants in PD genes as reported in our recent publications [12,13].

Healthy controls were matched for sex and age with PD patients and were negative for neurological disorders and for mutations/variants in PD genes.

#### MNI cohort used in case-control analysis

The Mediterranean Neurological Institute (MNI)-PD cohort included 804 independent and unrelated PD patients (501 males; 300 familiar and 504 sporadic cases), for which WES data are available. This cohort is part of the Parkinson’s disease Biobank of the IRCCS Neuromed and of the Institute of Genetics and Biophysics (CNR). All the subjects were of European ancestry and were evaluated by qualified neurologists of the Parkinson Centre of the IRCCS INM Neuromed from June 2015 to December 2017, and from June 2021 to December 2023, with a thorough protocol comprising neurological examination and evaluation of non-motor domains. Information about family history, demographic characteristics, anamnesis, and pharmacological therapy was also collected (the treatment of the PD groups consisted for the most part of a combination of levodopa and dopamine agonists). This cohort was used to identify genetic factors associated with either increased susceptibility or protection against PD.

#### Italian control cohort

282 neurological controls for which WES data are available. They were recruited by the same group of neurologists, among the patients’ wives/husbands, after having ascertained the lack of neurological pathologies and the absence of affected family members.

#### EUR-nfe cohort

404 healthy non-Finnish European population subjects from the 1000 Genomes Phase 3, data annotated in the GRCh38 assembly were downloaded from https://www.internationalgenome.org/, integrated snvindels version 2a 27022019 GRCh38 e.g. ftp://ftp.1000genomes.ebi.ac.uk/vol1/ftp/data_collections/1000_genomes_project/release/20190312_biallelic_SNV_and_INDEL/ALL.chr14.shapeit2_integrated_snvindels_v2a_27022019.GRCh38.phased.vcf.gz

#### Replication cohort

4586 PD patients from Parkinson Disease Genetic Consortium (PDGC) and 43989 controls from United Kingdom (UK) biobank, whose data were downloaded from the PDGC Variant browser (https://pdgenetics.shinyapps.io/VariantBrowser/).

### Generation of human induced Pluripotent Stem Cell (iPSC)

Peripheral Blood Mononuclear Cells (PBMC) were isolated from 15 mL of total blood using SepMate™ PBMC Isolation Tubes. Lymphoprep was used to generate the density gradient medium. PBMCs were collected from the Buffy coat using sterile Pasteur pipettes and cryopreserved in freezing medium (50% RPMI medium, 40% fatal bovine serum and 10% dimetilsolfossido). Human iPSCs were generated using the Epi5™ Episomal iPSC Reprogramming Kit, according to the manufacturer’s protocol for Reprogram StemPro™ CD34+ Cells, with appropriate optimizations. Day –3: PBMCs were tawed and plated in a 24-well plate in complete StemPro™-34 medium supplemented with cytokines (i.e., SCF 100 ng/mL, IL-3 20 ng/mL, IL-6 20 ng/mL and Flt3-ligand 100 ng/mL).

Day 0: 2×10^6^ T cells were transfected using the Amaxa nucleofector System. Transfected cells were plated onto Geltrex™ matrix-coated culture dishes and incubated overnight in complete StemPro™-34 medium containing cytokines (SCF (100 ng/mL), IL-3 (20 ng/mL), IL-6 (20 ng/mL) and Flt3-ligand (100 ng/mL)).

Day 1-8: medium was replaced with N2B27 Medium supplemented with 100 ng/mL bFGF and changed every other day.

Day 9: medium was replaced with mTeSR™ Medium, and cultures were monitored for the emergence of iPSC colonies. Medium was refreshed every other day.

Day 15-21: undifferentiated iPSC colonies were manually piked and transferred onto fresh Geltrex™ matrix-coated culture dishes for further expansion.

### Human iPSC culture

Human iPSCs were maintained and expanded in mTeSR medium for approximately 10 passages using mechanical dissociation of colonies. Cells were passaged at ∼70% of confluence. Briefly, colonies were visualized under an optical microscope, and morphologically optimal colonies lacking differentiation areas were manually subdivided into small fragments (typically, each colony was divided into approximately nine pieces). Approximately 20/30 fragments derived from different colonies were transferred into a new well for continued expansion.

After ten passages, human iPSCs were maintained and expanded into StemMACS™ iPS-Brew XF medium. Cell expansion at this stage was performed using chemical dissociation with phosphate-buffered saline (PBS) supplemented with 0.5 mM Ethylenediaminetetraacetic acid (EDTA).

Human iPSCs were cryopreserved in CS10 Cryopreservation Media (SigmaAldrich) according to the manufacturer’s instructions and stored in liquid nitrogen.

Karyotype analysis was performed to evaluate the chromosomal integrity of selected iPSC clones. G-banding analysis was conducted by MeriGen laboratory SPA (Italy). More than 10 metaphases were analysed for each clone. Karyotype assessment was performed after approximately 10 passages, when clones had fully lost episomal vectors.

### RNA Expression studies

Total RNA from hiPSC was isolated from samples using Trizol kit (Invitrogen) protocol. Five µgs of total RNA were digested with TURBO™ DNase (RNase free) kit (Thermo Fisher Scientific, Waltham, MA, USA) to eliminate genomic DNA contamination. Two micrograms of DNA-free RNA were reverse transcribed with the Superscript III-First strand kit (Thermo Fisher Scientific, Waltham, MA, USA). Quantitative PCR (qPCR) reactions were performed in triplicate, using gene specific primers **(Table S.1)** and ITaq Universal Sybr Green Supermix (Bio-Rad, Hercules, CA, USA) following the manufacturer’s directions. Results were normalized to the expression of the beta actin (ACTB) gene. Standard deviation was calculated by using data of three different experiments.

### Digital droplets PCR (ddPCR)

ddPCR was used to test the expression of lowly expressed genes in mesencephalic dopaminergic (mesDA) neurons. The absolute number of target molecules is calculated using Poisson statistical analysis and expressed as copies per microliter (copies/µL), without the need for calibration curves or external standards. In this study, droplets were generated with Droplet Generation Oil for Probes (Bio-Rad) by using QX200/QX100 droplet generator. The PCR mix was prepared with the ddPCR™ Supermix for Probes according to the manufacturer protocol (Bio-Rad) with a final reaction volume of 22 µL. The amplification of the molecular target was conducted with the C1000 Touch Thermal Cycler. All the experiments were performed in triplicate and analysed with QX200 Droplet Reader with QX Manager software. The list of the probes adopted for expression studies is reported in **Table S.1**.

### Immunofluorescence (IF) staining of human iPSC cell lines

The human iPSCs were grown in 24 wells plate using complete medium for 24/48 h. Colonies were fixed with paraformaldehyde 4% and stained with anti-SOX2, anti-OCT4, anti-NANOG, and anti-CRIPTO antibodies. Briefly, incubation with blocking buffer (1×PBS/0.1% triton/5% donkey serum) was performed for 2 hours at room temperature (RT) on shaker. Primary and then secondary antibody reactions (1×PBS/0.1% triton/antibody) were performed at 4°C overnight and at RT for 1 hours, respectively. Anti-SOX2 (MAB2018, R&D System, UK, 1:200), anti-OCT4 (Sc-5279, Santa Cruz, UK, 1:100), anti-NANOG (8822, Cell Signaling, Danvers, Massachusetts, USA, 1:400), anti-CRIPTO (2775, Cell Signaling, Danvers, Massachusetts, USA, 1:200), anti-FOXA2 (sc-101060, SantaCruz 1:2000), anti-MAP2 (AB5622, Merck Millipore 1:200), anti-TH (AB152, Merck Millipore), anti-LMX1 (AB10533, Merck Millipore 1:1000), Donkey anti-Mouse IgG (H+L) Highly Cross-Adsorbed Secondary Antibody, Alexa Fluor™ 568 (A10037, Thermo Fisher Scientific CA USA, 1:250) and Donkey anti-Rabbit IgG (H+L) Alexa Fluor™ 488 (A-21206, Thermo Fisher Scientific CA USA, 1:250) secondary antibodies were used. Hoechst H3570 (Thermo Fisher Scientific CA USA, 1:1000) was used to visualize the nuclei. Slides were visualized with Nikon Confocal Microscope A1R.

Quantification of TH-positive cells was performed on images acquired from fixed samples immunostained for TH, with nuclei counterstained with DAPI. Image analysis was performed using Image J 1.46r software (https://imagej.nih.gov/ij/). For each analyzed field, TH-positive cells were counted and normalized to the area (the area was 0.284 mm^2^ for each acquired image).

### Generation of human Ventral midbrain dopaminergic neurons

Human Ventral midbrain dopaminergic neurons were generated following previously described protocols by Nolbrant S. and colleagues [15].

Briefly, human iPSCs were dissociated with EDTA 0.5mM, collected and counted. A total of 2×10^4^ cells were plated into 24-well plates precoated with laminin-111 (Lam-111; 1 µg/cm2). Cells were differentiated in N2 medium (50% Neurobasal and 50% D-MEM F12 supplemented with N2) containing SB431542 (10 µM), Noggin (100 ng/ml), Shh-C24II (300 ng/ml), and CHIR99021 (0,9 µM) for 9 days. Medium was changed on days 2, 4 and 7. On day 9, medium was replaced with N2 medium supplemented with FGF8b (100 ng/ml).

On day 11, cells were dissociated and replated. A total of 1.6×10^6^ cells were seeded into 24-well plate precoated with laminin-111 (Lam-111; 1 µg/cm2). Cells were further differentiated using B27 medium (Neurobasal supplement with B27) containing BDNF (20 ng/ml), AA (0.2 mM), and FGF8b (100 ng/ml) until day 16. Medium was replaced on day 12 and day 14.

On day 16, cells were replated for terminal in vitro differentiation. A total of 3.1×10^5^ cells were seeded into 24-well plate pre-coated with laminin-111 (2 µg/cm²). Cells maintained in B27 medium supplemented with BDNF (20 ng/ml), AA (0.2 mM), GDNF (10 ng/ml), db-cAMP (500 µM), and DAPT (1 µM) for approximately 3 weeks. Complete medium changes were performed every 3 days until day 25, followed by half-medium changes every 3 days thereafter.

### Electrophysiological recordings

Patch-clamp recordings were performed at 45/50 days of differentiation in hIPSC-derived dopaminergic neurons by using Multiclamp 700B amplifier and Axon Digidata 1550A (Molecular Devices, CA). Post-synaptic currents were recorded at a 10 KHz sampling rate (2KHz filter) and analyzed by PClamp software (Molecular Devices). Action potentials (APs) were evoked by a current clamp step protocol (from –40 pA to 100 pA, delta level +20 pA, 500 ms in duration, 20 KHz sampling rate). The analysis of AP parameters was performed evoking a single AP by a short step protocol (150-200 pA, 30 ms duration). For details, see Martinello et al., 2025 [44]. Normal extracellular solution (NES) contained (in mM) NaCl 140, HEPES 10, KCl 2.8, CaCl2 2, MgCl2 2, and glucose 10, pH 7.4 with NaOH. The intracellular solution contained (in mM) KCl 140, HEPES 10, MgCl2 2, MgATP 2, EGTA 0.5, pH 7.4 with KOH.

### Sample preparation for lipidomic and proteomic analysis

hiPSC-derived dopaminergic neurons were grown in quadruplicate in 24 wells plate for 45 days. For lipidomic analysis, 3×10^5^ cells were processed using 1 mL of 75:15 IPA/H2O solution, after the addition of 100 μL of CH3OH 5% deuterated standard (Splash Lipidomix®). Then the samples were vortexed for 30 s, sonicated for 2 min, vortexed again for 30 s and then they were incubated for 30 min at 4 °C, under gentle, constant shaking. Subsequently, samples were rested on ice for an additional 30 min. To remove debris and other impurities, the samples were centrifuged for 10 min at 3500×g at 4 °C. 1 mL of supernatant was collected and dried using a SpeedVac centrifuge (Labogene). The dried samples were reconstituted in 100 μL of CH3OH containing the internal standard CUDA (12.5 ng/mL). For proteomic analysis, 1×10^5^ cells were lysed with RIPA buffer and sonicated. Proteins were then precipitated with cold acetone and resuspended. Proteins were then reduced in 25 µL of 100 mM NH4HCO3 with 2.5 μL of 200 mM DTT (Merck) at 60 °C for 45 min and next alkylated with 10 μL 200 mM iodoacetamide (Merck) for 1 h at RT in dark conditions. Iodoacetamide excess was removed by the addition of 200 mM DTT. Proteins were then digested with trypsin. The digests were dried by Speed Vacuum and then desalted [45].

### Lipidomic analysis

For lipidomic analysis, reconstituted samples were analysed with a Vanquish UHPLC system (Thermo Scientific, Rodano, Italy) coupled with an Orbitrap Q-Exactive Plus (Thermo Scientific, Rodano, Italy). Lipid separation was performed using a reversed-phase column (Hypersil Gold™ 150 × 2.1 mm, particle size 1.9 µm) maintained at 45 °C with a flow rate of 0.260 mL/min. Mobile phase A for ESI mode positive consisted of 60:40 (v/v) acetonitrile/water with ammonium formate (10 mmol) and 0.1% formic acid, while mobile phase B was 90:10 isopropanol/acetonitrile (v/v) with ammonium formate (10 mmol) and 0.1% formic acid, while in the negative ESI mode, the organic solvents for both mobile phases were the same as in the positive with the exception of using ammonium acetate (10 mmol) as a mobile phase modifier. The gradient used was as follows: 0–2 min from 30 to 43% B, 2–2.1 min from 43 to 55% B, 2.1–12 min from 55 to 65% B, 12–18 min at 65% to 85% B, 18–20 min at 85% to 100% B; 100% B was held for 5 min, and then the column was allowed to equilibrate to 30% B for another 5 min. The total running time was 30 min. Mass spectrometry analysis was performed in both positive ion (at 3.5 kV) and negative ion (2.8 kV) modes. Data were collected in a data-dependent top 10 scan mode (ddMS2). MS full-scan survey spectra (mass range m/z 80–1200) were acquired with a resolution of R = 70,000 and target AGC of 1 × 106. MS/MS fragmentation was performed using high energy c-trap dissociation (HCD) with R = 17,500 resolution and 1 × 105 AGC target. The step normalised collision energy (NCE) was set to 15, 30 and 45. The injection volume was 3 µL. For accurate mass-based analysis, regular Lockmass and interrun calibrations were used. An exclusion list for background ions was generated by testing the same procedural sample for both positive and negative ESI modes22. Quality control was ensured by analysing pooled samples before, at the beginning and at the end of the batches; using blanks to check for residual interference; and using internal standards, directly in plasma or cell samples, which include a series of analyte classes at levels appropriate for the plasma (Avanti SPLASH Lipidomix) and an internal standard (CUDA) prior to liquid chromatography-mass spectrometry (LC–MS) analysis.

Raw data acquired from lipidomic untargeted analysis were processed with MSDIAL software (Yokohama City, Kanagawa, Japan), version 4.24. Peaks were detected, MS2 data were deconvoluted, compounds were identified, and peaks were aligned across all samples. For quantification, the peak areas for the different molecular species detected were normalised using the deuterated internal standard for each lipid class. To obtain an estimated concentration expressed in nmol/mL (plasma), the normalised areas were multiplied by the concentration of the internal standard. An in-house library of standards was also used for lipid identification.

### Proteomic analysis

Digested peptides were analysed on an Ultimate 3000 RSLC nano coupled directly to an Orbitrap Exploris 480 with a High-Field Asymmetric Waveform Ion Mobility Spectrometry System (FAIMS) (all Thermo Fisher Scientific). Samples were injected onto a reversed-phase C18 column (15 cm × 75 µm i.d., Thermo Fisher Scientific) and eluted with a gradient of 6% to 95% mobile phase B over 41 min by applying a flow rate of 500 nL/min, followed by an equilibration with 6% mobile phase B for 1 min. The acquisition time of one sample was 41 min and the total recording of the MS spectra was carried out in positive resolution with a high voltage of 2500 V and the FAIMS interface in standard resolution, with a CV of −45V. The acquisition was performed in data-independent mode (DIA): precursor mass range was set between 400 and 900, isolation window of 8 m/z, window overlap of 1 m/z, HCD collision energy of 27%, orbitrap resolution of 30,000 and RF Lens at 50%. The normalised AGC target was set to 1000, the maximum injection time was 25 ms, and microscan was 1. For DIA data processing, DIA-NN (version 1.8.1) was used: the identification was performed with “library-free search” and “deep learning-based spectra, RTs and IMs prediction” enabled. Enzyme was set to Trypsin/P, precursors of charge state 1–4, peptide lengths 7–30 and precursor m/z 400–900 were considered with maximum two missed cleavages. Carbamidomethylation on C was set as fixed modification and Oxidation on M was set as variable modification, using a maximum of two variable modifications per peptide. FDR was set to 1%.

The mass spectrometry proteomics data have been deposited to the ProteomeXchange Consortium via the PRIDE partner repository with the dataset identifier PXD074629.

### Statistical analysis

Statistical analyses of lipidomic and proteomic samples were performed with MetaboAnalyst 6.0 (www.metaboanalyst.org). Statistical Analysis [single factor] function was used to build volcano plots, principal component analysis (PCA), partial least squares-discriminant analysis (PLS-DA) and heatmaps. Pathway and Enrichment analysis functions were performed to integrate pathway enrichment analysis and pathway topology analysis, starting from significant proteins found in statistics. GraphPad Prism 8 was used for unpaired Student’s T-test, ONE-way ANOVA and multiple comparison with post hoc correction statistical analysis.

Bioinformatic analysis was carried out using DAVID software (https://david.ncifcrf.gov/tools.jsp) and Ingenuity Pathway Analysis (Qiagen, Hilden, Germany).

### Association analysis

Principal Component Analysis (PCA) was performed with PLINK software and was used to characterize the genetic diversity of the study sample (PD_MNI, controls (CNT)_MNI, EUR_nfe) [46]. The analysis was carried out by using common variants (Minor allele frequency MAF > 0.01), PC1 and PC2 were found to contribute to a variance of 25% among samples.

To identify the genetic contribution given by common variants (MAF > 0.01) we adopted a logistic regression under an additive genetic model trough PLINK2 software, by adjusting for sex and the 10 principal components. p-values were adjusted for Bonferroni multiple testing correction.

### Expression studies in human brain regions

The Genotype-Tissue Expression (GTEx) portal (https://www.gtexportal.org/home/) was accessed to obtain gene expression data of the identified SNPs. The analysis was performed on all available adult human brain regions (amygdala, anterior cingulate cortex BA24, caudate nucleus, putamen, substantia nigra, cerebellar hemispheres, cerebellum, cerebral cortex, frontal cortex BA9, hippocampus, hypothalamus, nucleus accumbens).

## Results

### Study cohort and generation of human induced pluripotent stem cell lines

To investigate the synergistic effects of multiple variants or mutations in PD-associated genes we selected six PD patients carrying single or multiple highly penetrant variants in 11 PD candidate genes (*AIMP2, ANKK1, HMOX2, KIF21B, LRRK2, RHOT2, SLC6A3, TMEM175, TOMM22, TVP23A* and *ZSCAN21*) [12,47], as well as three healthy subjects (HS) without neurological diseases, at the time of sample collection, and negative for mutations/variants in PD-related genes.

Specifically, *KIF21B* (PD1) encodes a kinesin motor protein involved in microtubule dynamic and mitochondrial transport along neuronal axons [48]; *ZSCAN21* (PD2) is a transcription factor implicated in the regulation of *SNCA* expression [49–51]; *TMEM175* (PD3 and PD6) encodes a lysosomal K^+^ channel essential for lysosome physiology, autophagy and α-Synuclein clearance [52]; *HMOX2 (*PD4*), RHOT2* and *TOMM22* (PD5) contribute to mitochondrial homeostasis [53–56]; *AIMP2* (PD6) is a Parkin substrate associated with activation of the *parthanatos* cell death pathway [57–59]*; SLC6A3* (PD4) encodes the Dopamine Transporter 1 (DAT) whose dysfunction underlies *SLC6A3*-related dopamine transporter deficiency syndrome [60]; and *ANKK1* (PD2) encodes a serine/threonine kinase of the receptor-interacting protein family, which plays key roles in the regulation of cell survival, death, and differentiation [61,62]. Finally, *TVP23A* (PD3) is a recently described gene (Gene ID: 780776) implicated in intracellular vesicular transport [12,13].

Demographic and clinical characteristics of all participants are summarized in **Table 1**, as well as the complete list of gene variants is reported in **Table S2**. The PD cohort included four sporadic PD patients (three males and one female) and two familial PD patients (both males), with age at symptom onset (AAO) ranging from 44 to 59 years. Except for patient PD3, who presented with a moderate clinical phenotype, the remaining patients (PD1, PD2, PD4, PD5, and PD6) exhibited severe disease manifestations affecting motor (MDS-UPDRS and HY scores), non-motor (NMS scales), and cognitive (MoCA score) domains (**Table 1**).

**Table 1.** Demographic and clinical characteristics of the study cohort from which hiPSCs were generated.

| ID_iPSCs | Sex | Age | AAO | Disease Duration | UPDRS | PD form | Phenotype | LID | MoCA | NMS | Mutated gene |
| --- | --- | --- | --- | --- | --- | --- | --- | --- | --- | --- | --- |
| HS1 | M | 61 | - | - | - | - | - | - | - | - | - |
| HS2 | F | 61 | - | - | - | - | - | - | - | - | - |
| HS3 | M | 61 | - | - | - | - | - | - | - | - | - |
| PD1 | F | 66 | 59 | 7 | 59 | Sporadic | Mixed | No | 15 | 200 | *KIF21B* |
| PD2 | M | 69 | 58 | 11 | 64 | Sporadic | Mixed | Yes | 10 | 200 | *ZSCAN21, ANKK1* |
| PD3 | M | 55 | 44 | 11 | 30 | Sporadic | Bradykinetic-Rigid | Yes | 30 | 32 | *TMEM175, TVP23A* |
| PD4 | M | 74 | 53 | 21 | 35 | Sporadic | Tremor-dominant | Yes | 28 | 116 | *SLC6A3, HMOX2* |
| PD5 | M | 61 | 48 | 13 | 30 | Familial | Bradykinetic-Rigid | Yes | 24 | 82 | *LRRK2, RHOT2, TOMM22* |
| PD6 | M | 76 | 54 | 22 | 56 | Familial | Mixed | No | 19 | 140 | *TMEM175, AIMP2* |

AAO: Age at Onset; UPDRS: Unified Parkinson's Disease Rating Scale; LID: Levo-Dopa induced Dyskinesia; MoCA: Montreal Cognitive Assessment; NMS: Total score of Non-Motor Symptoms; M: Male; F: Female; Age and AAO were reported in terms of years.

Human iPSC lines were generated by reprogramming PBMCs using the Epi5™ Episomal iPSC Reprogramming Kit according to the manufacturer’s instructions, with minor modifications (detailed in the Materials and Methods section). Briefly, PBMCs were cultured for three days in StemPro™-34 medium supplemented with cytokines (i.e., SCF 100 ng/mL, IL-3 20 ng/mL, IL-6 20 ng/mL and Flt3-ligand 100 ng/mL). Non-adherent-PBMCs were nucleofected with episomal vectors encoding Oct4, Sox2, Klf4, L-Myc, and Lin28 to induce pluripotency. Pluripotency stabilization was achieved by supplementation with β-Fgf, and successful induction was confirmed by Alkaline phosphatase staining (data not shown). Individual colonies were mechanically dissociated and expanded to ensure genomic stability. We obtained about 5/6 clones for lines apart from PD1 for which only 3 clones were collected.

A schematic overview of the reprogramming workflow is shown in **Supplementary Fig.1a**, and detailed protocols are provided in Materials and Methods.

A comprehensive characterization protocol was applied to ensure the high-quality standard of all the generated hiPSC lines. Once established, the hiPSC lines were expanded for at least 5 passages prior to characterization, which included: (1) assessment of colony morphology resembling human pluripotent stem cells (**Supplementary Fig.1b**); (2) confirmation of episomal vector loss via PCR using primers against episomal genes (ORI-P); (3) evaluation of pluripotency and self-renewal markers at both the RNA (qPCR) and protein (immunofluorescence) levels; (4) karyotype analysis to detect possible chromosomal abnormalities; and (5) assessment of neural differentiation potential. Donor cells were screened for the absence of viral infections (HIV, HCV, HBV) and mycoplasma contamination, ensuring that no confounding factors affected cellular integrity.

Episomal vector clearance was monitored using end-point PCR. Clones derived from different individuals showed variable episomal loss rates; on average, approximately ten passages were required to obtain integration-free hiPSC lines **(Supplementary Fig.1c)**. No significant differences were observed between iPSCs derived from patients and those from healthy subjects **(Supplementary Fig.1c)**.

To confirm pluripotency, we analyzed markers of pluripotency and self-renewal at the RNA and protein levels. Expression of pluripotency-associated genes (i.e. *NANOG, SOX2, CRIPTO, OCT4, KLF4*) was evaluated by qPCR by using primers designed to recognize endogenous transcripts. All hiPSC lines exhibited a robust expression of *NANOG, SOX2, OCT4, CRIPTO[63]*, accompanied by reduced *KLF4* expression relative to the parental PBMCs. A well-characterized pluripotent stem cell (PSC) line was used as an internal control for reprogrammed PD and HS clones [64] (**Fig.1 and Supplementary Fig. 2**). Although expression levels varied among individual clones, all reprogrammed lines exhibited comparable levels of pluripotency-associated markers and successfully passed quality control criteria based on human PSC standards (**Fig. 1 and Supplementary Fig. 2**). These results show robust activation of the endogenous pluripotency transcriptional network and confirm efficient reprogramming from a differentiated to an undifferentiated state. Consistently, immunofluorescence analyses confirmed uniform expression of SOX2, OCT4, NANOG, and CRIPTO across all reprogrammed lines (**Fig. 1 and Supplementary Fig. 2**).

**Figure 1:**
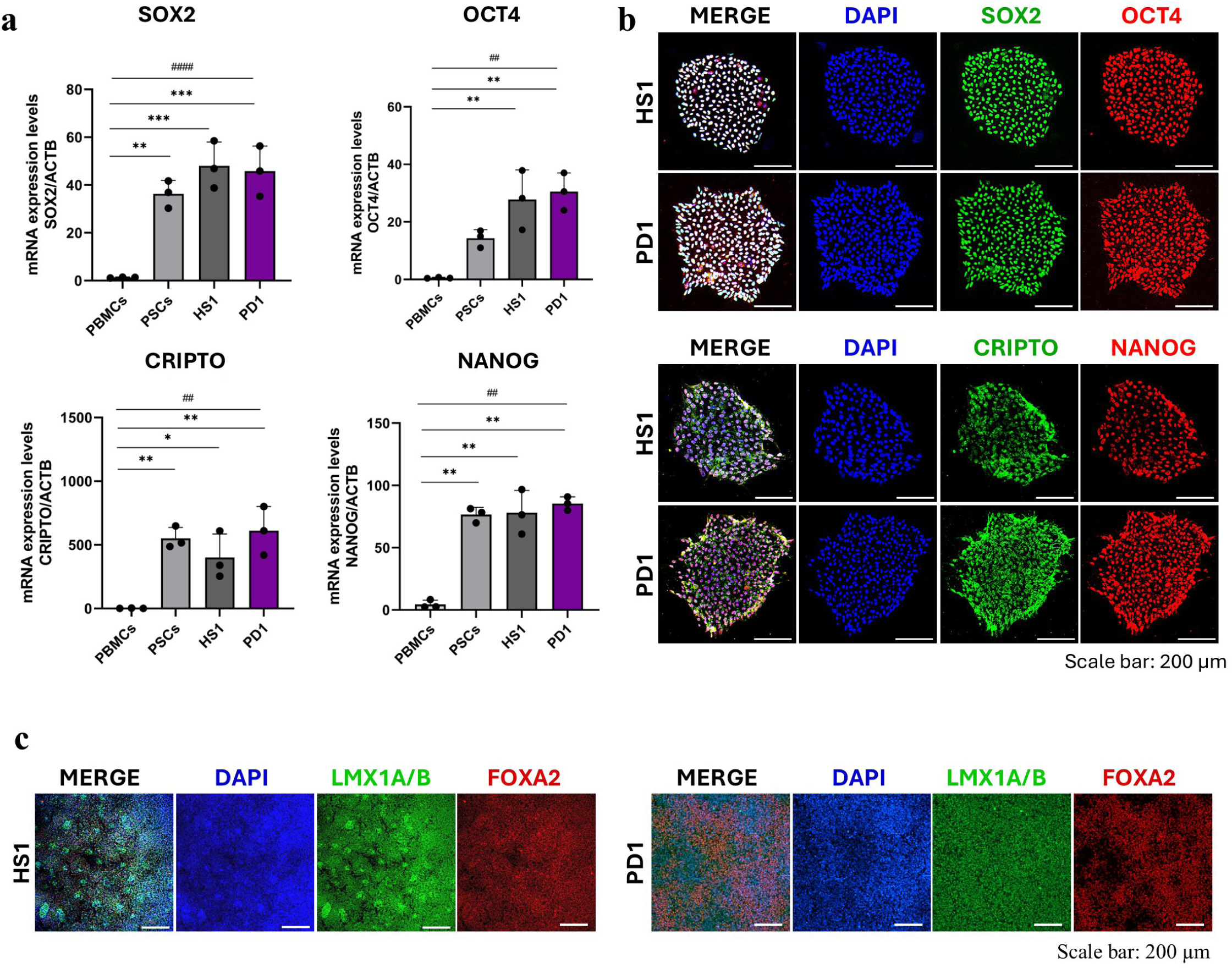
Generated hiPSCs displayed the presence of pluripotency markers and differentiated in neuronal progenitor cells. **a)** Real-time PCR indicate increased mRNA expression level of pluripotency genes *SOX3*, *OCT4*, *CRIPTO* and *NANOG* in HS1 and PD1 reprogrammed cell lines. All the values were expressed as arbitrary unit and normalised on *ACTB* (B-Actin). PSCs were used as control of the reprogramming protocol. Data are represented as Mean±SD from three independent experiments. # shows the significance calculated with one-way ANOVA to assess different expression value among the groups. #p<0.05, ##p<0.01, ### p<0.001, #### p<0.0001. * indicates comparison respect to PBMCs evaluated by multiple comparison’s test after Dunnett’s post hoc correction. Significance was set at p<0.05. * p<0.05, ** p<0.01, *** p<0.001, **** p<0.0001. **b)** Immunofluorescence staining was showed on hiPSCs representative single colony image of HS1 and PD1 stained with DAPI (in blue), SOX2 and CRIPTO in green, OCT4 and NANOG in red. **c)** IF analysis confirmed the presence of ventral midbrain (VM) progenitor cells positive for LMX1A/B (in green) and FOXA2 (in red) at day-16; nuclei were stained with DAPI. Scale bar: 200µm

**Figure 2:**
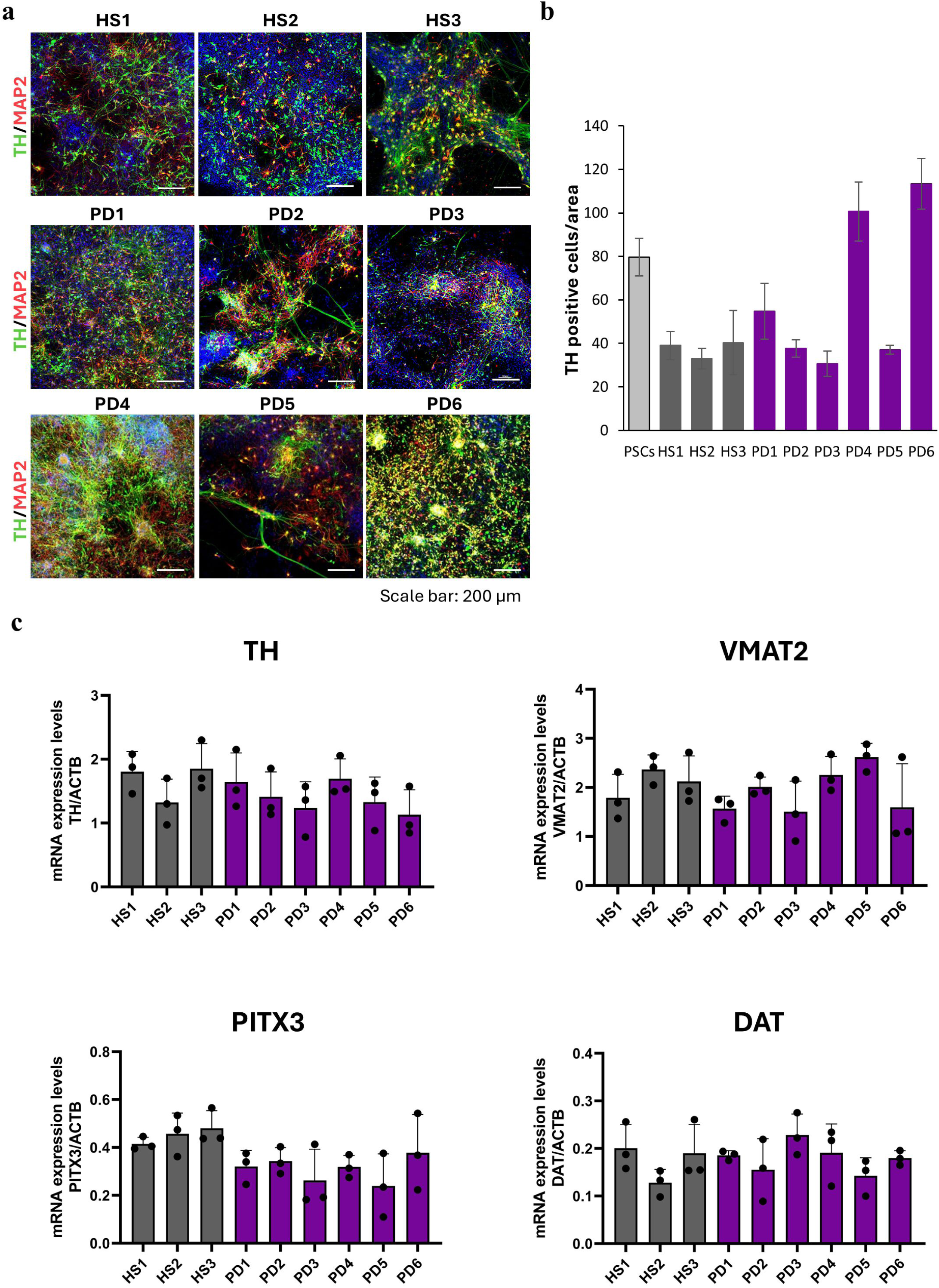
mesDA neurons-derived from HS and PD patients displayed presence of mature dopaminergic markers at 45 days of differentiation. **a)** Immunofluorescence analysis detected positive neurons for TH (green) and MAP2 (red) at 45 days of differentiation. **b)** Graphical representation of TH count of positive cells per area showing a similar number of neurons positive for TH in HS and PD cell lines. One-way ANOVA was adopted to assess significant differences among the groups. **c)** Real-time PCR analysis confirmed expression at mRNA level of dopaminergic mature markers including *TH*, *DAT*, *VMAT2* and *PITX3*. Data are represented as MEAN±SD from three independent experiments. One-way ANOVA was adopted to assess significant differences among the groups.

To assess genomic integrity, we performed karyotype analyses on twenty-seven independent clones (three clones per line; ten metaphases per clone). Most clones displayed a normal karyotype, with chromosomal abnormalities detected in only two clones (Monosomy X and 46,XY,add(18)(p11.3)) (**Supplementary Fig.1d**). Overall, these results indicate a high level of genomic stability across the reprogrammed lines, consistent with the use of mechanical dissociation during colony selection and expansion to minimize reprogramming-induced genomic alterations [65]. Importantly, the frequency of chromosomal abnormalities did not differ between PD– and control-derived hiPSC lines.

### Differentiation of PD iPSC lines into mature dopaminergic neurons

In vitro modelling of Parkinson’s disease requires the fine and efficient generation of mesencephalic dopaminergic (mesDA) neurons, which arise from a distinct population of progenitors located in the floor-plate of the ventral midbrain (VM), characterized by LMX1A+/FOXA2+ expression [15]. The in vitro derivation of these neurons remains challenging due to the difficulty of recapitulating the floor plate microenvironmental and the dependence on iPSC quality. Nevertheless, several differentiation protocols have been established to overcome these limitations.

In this study, we adopted the protocol described by Nolbrant and colleagues [15], which enables efficient neural induction through dual SMAD inhibition, followed by precise caudalization using a glycogen synthase kinase 3 inhibitor (GSK3i) and ventralization/floor-plate induction with Sonic Hedgehog (SHH-C24II). This approach promotes the generation of FOXA2+/LMX1+/PAX6− VM floor-plate progenitors. The timeline and factors used for differentiation are detailed in the Materials and Methods section.

The progression of differentiation toward C neurons was assessed at day-16 and day-45. By day-16, cultures robustly generated late caudal ventral midbrain (VM) progenitors co-expressing LMX1 and FOXA transcription factors (**Fig. 1** and **Supplementary Fig. 3**). iPSC-derived progenitors displayed a largely consistent LMX1 expression pattern relative to PSCs, with only minor intercellular variability. Importantly, no significant differences were observed between progenitors derived from PD patients and healthy controls, indicating comparable differentiation efficiency across all lines. By day 45, the presence of mature DA neurons was assessed by immunofluorescence analysis for Tyrosine Hydroxylase (TH) and MAP2 **(Fig. 2a, b; Supplementary Fig. 4).** Quantification of TH⁺ neurons revealed robust dopaminergic differentiation with no significant differences between PD-derived and control lines. In parallel, quantitative PCR analysis of post-mitotic dopaminergic markers, including *TH*, *VMAT2*, *PITX3*, and *DAT*, further confirmed the acquisition of a mature dopaminergic neuronal identity in culture (**Fig. 2C**).

### PD-derived dopaminergic neurons with different genetic backgrounds exhibit functional alterations of both excitability and glutamatergic synaptic transmission

To describe the possible functional differences of mesencephalic dopaminergic (mesDA) cultures derived from HS and PD patients, we recorded action potentials (APs) and excitatory post-synaptic currents from HS– and PD-derived neurons at days 45-48. We found significantly smaller capacitance in PD1-, PD4-, PD5-, and PD6-derived neurons compared to HS, but similar resting membrane potential **(Fig. 3a and b)**. Moreover, all the PD-derived cultures exhibited APs **(Fig. 3c)**. Eliciting APs by a current clamp step protocol, we observed a reduced number of APs in PD1, PD4, PD5, and PD6 compared to HS (**Fig. 3d, e**). In PD1, PD4, PD5, and PD6, we also observed smaller AP amplitudes, higher AP thresholds, and reduced half-width and after-hyperpolarization values **(Fig. 3f-k)**. These data suggest a lower maturation of PD-derived neurons compared to HS. Interestingly, we observed an increase in AP numbers in PD3 compared to HS cells, while all other AP parameters were similar to those of HS cells.

**Figure 3:**
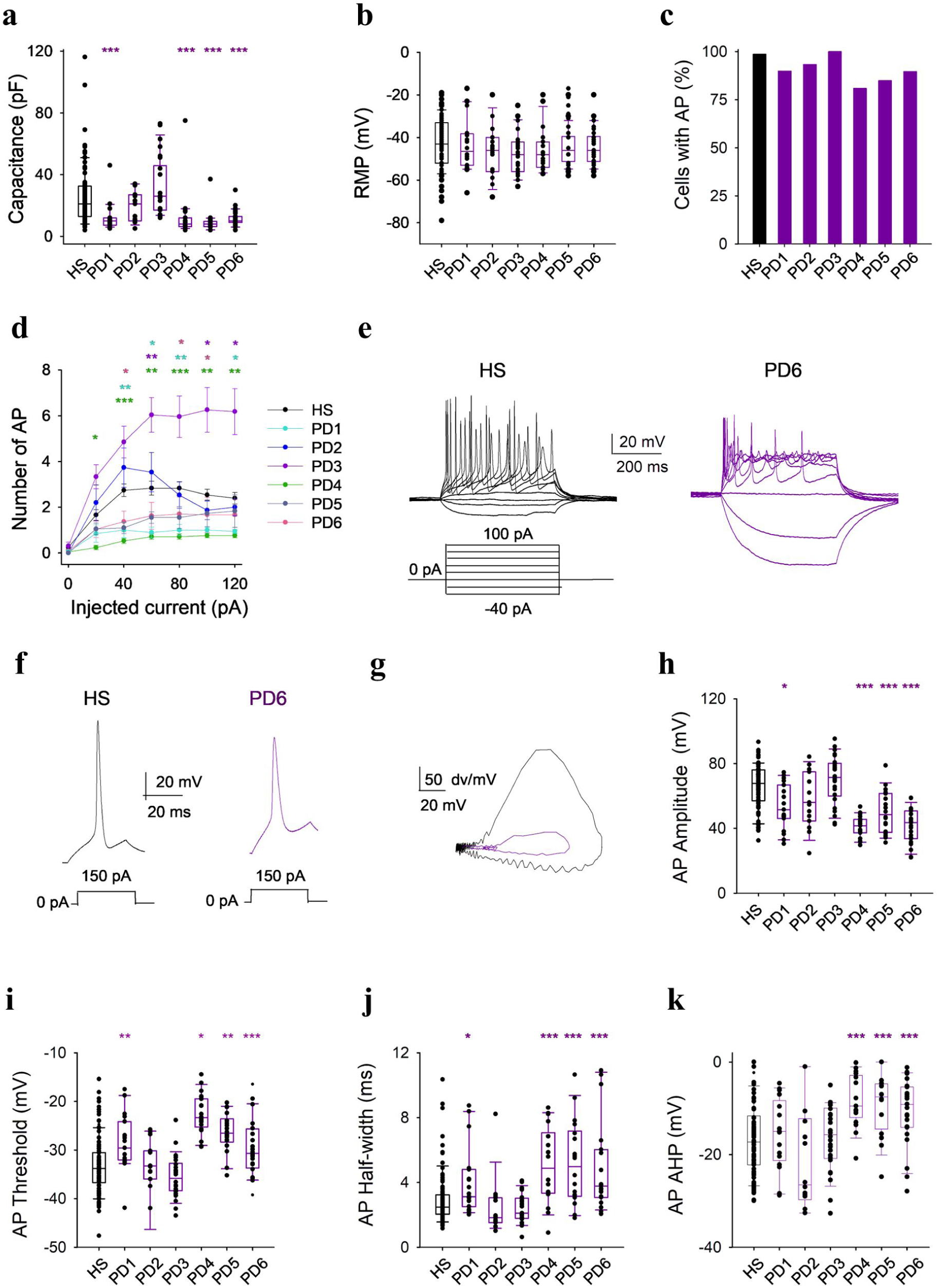
Parkinson-associated mutations cause alterations in neuronal excitability. **a)** Box-plot illustrating the membrane capacitance values obtained from HS– and PD-derived dopaminergic neurons; 4 out of 6 different mutations are associated with a significant decrease in the capacitance when compared with the HS (***p<0.0001; n=89, 20, 15, 26, 20, 20 and 31, for HS and PD1 to PD6, respectively); **b)** Resting membrane potentials recorded from same cells as in **a; c)** Bar graph illustrating the percentage of cells (same as in A and B) exhibiting action potentials; **d)** Input-output curves showing the difference in number of action potentials (APs) elicited by the CC protocol in HS– and PD–derived dopaminergic neurons (same cells as in A; ***p<0.0001; **p<0.001; * p<0.05 One Way ANOVA, versus HS); **e)** Typical traces obtained from dopaminergic neurons during CC step protocol (from –40 pA to 120 pA, 500 ms step duration) from a HS (black traces) and a PD6-derived neuron (violet trace); **f)** Typical traces obtained from dopaminergic neurons during CC single step protocol (150 pA, 30 ms) from a HS (black traces) and a PD6-derived neuron (violet trace); **g)** Superimposed phase plane plots of the AP traces shown in F; **h)-k)** Box-plots representing the AP functional parameters measured from the same cells as in A (***p<0.0001; **p<0.001; * p<0.05 One Way ANOVA, versus HS).

Moreover, we observed a reduced number of synaptically active cells in PD4 and PD5, which carry pathogenic mutations in DAT and LRRK2, as well as in PD6 **(Fig. 4a)**. Furthermore, excitatory post-synaptic currents (EPSCS) recorded from PD4-derived cells displayed a smaller amplitude and a faster decay-time, causing a reduction of EPSC mean charge **(Fig. 4a, e and f)**. No change was observed in the frequency of EPSCs in any cell cultures **(Fig. 4g and h)**. Overall, these results revealed altered cell capacitance and action potential properties among PD-derived neurons, suggesting functional heterogeneity within the patient group.

**Figure 4:**
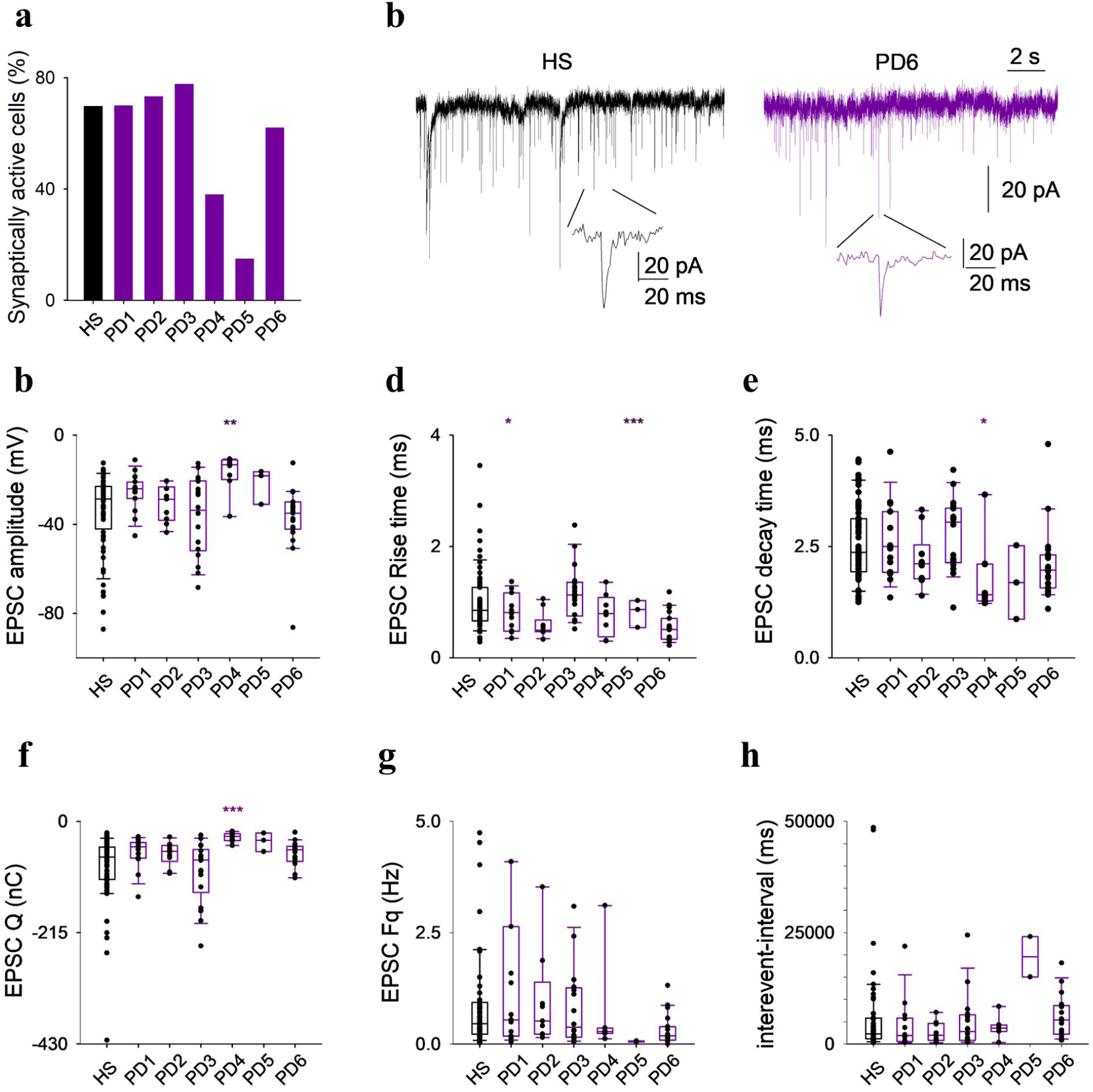
The glutamatergic synaptic transmission recorded from PD4– and PD5-derived neurons exhibits functional alterations. **a)** Percentage of HS– and PD– derived dopaminergic neurons showing excitatory post-synaptic currents (EPSCs; n=64, 15, 10, 18, 8, 3 and 19, for HS and PD1 to PD6, respectively); **b)** typical recordings of EPSCs obtained from dopaminergic neurons in a HS (black traces) and a PD6-derived neuron (violet trace); **c-f)** Box-plots representing the EPSCs functional parameters measured from the same cells as in A; **g**-**h)** box plots representing the EPSCs frequency and inter-event interval (n=59, 15, 9, 16, 7, 2 and 19 for HS and PD1 to PD6, respectively), (***p<0.0001; **p<0.001; * p<0.05 One Way ANOVA, versus HS).

### PD patient-derived dopaminergic neurons revealed specific alterations of lipidomic profile

To explore the lipidomic signature in PD cells, we collected iPSC-derived dopaminergic neurons at day-45 from 6 PD patients (PD1, PD2, PD3, PD4, PD5 and PD6) and 5 healthy subjects (3 lines were generated in this study and two lines were from external biobanks (see materials and methods)). Analyses were carried out through mass spectrometry in four independent biological replicates.

We identified and quantified 1265 expressed lipid species belonging to 40 lipid classes. Hierarchical heat map and Partial Least Squares Discriminant Analysis (PLS-DA) clearly discriminated the neuronal cells of PD patients with respect to healthy subjects (**Fig. 5a,b**). We identified 141 significant up-regulated (Fold Change (FC) ≥ 1.3; p value (p) ≤ 0.01) and 111 significant down regulated (FC ≤ 0.70; p ≤ 0.01) lipids in patient-derived neurons compared to controls (**Fig. 5c**). The complete dataset of lipids identified in this study is reported in **Table S3**. In the patient-derived neurons we observed increased levels of several lipid classes such as Acylcarnitine (CAR), Fatty Acids (FA), Ceramide (Cer), Hexosylceramide (HexCer), Sulfoglycosphingolipid (SHexCer), Glycerophospholipids (Phosphatidylcholine (PC), Ether-linked Phosphatidylcholine (PC-O), Phosphatidylethanolamine (PE)) and Glycerolipids like Diacylglycerols (DG) **(Fig. 5e)**. In PD cell lines, we also observed a highly significant reduction of the ganglioside GM3, which is the most abundant ganglioside in neural plasma membranes essential for signaling processes and compartmentalization of neurotransmitter components [66,67] (**Fig. 5e**).

**Figure 5:**
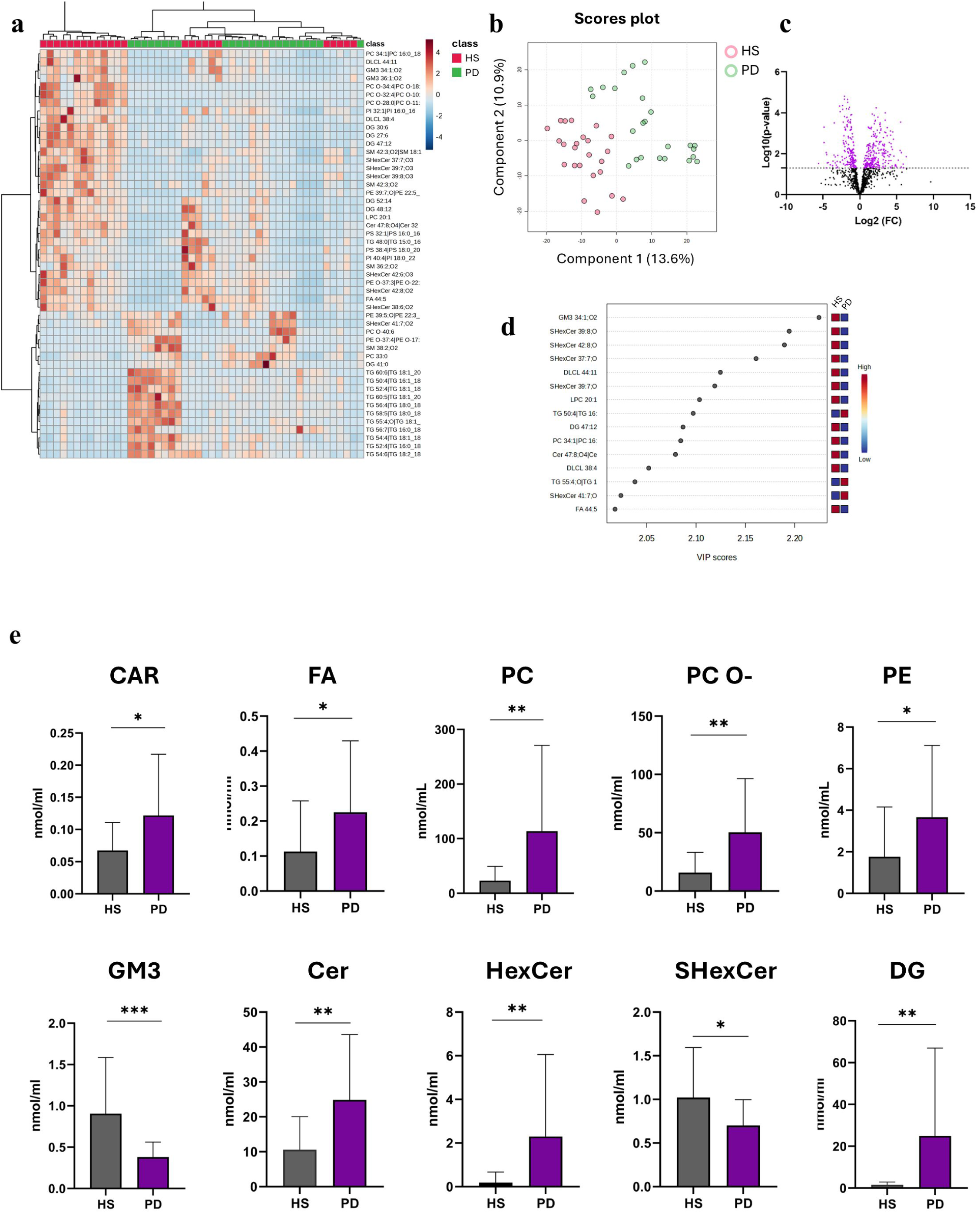
Lipidomic Analysis revealed altered lipid class in PD-derived mesDA neurons. **a-b**) Heatmap and Partial Least Square Analysis (PLSA) showed a defined clusterization and definition between the mesDA neurons derived from PD patients and controls. **c)** Volcano plot showing the distribution of significantly altered lipid species. **d)** VIP score analysis showed the most significantly altered lipid species when comparing PD-derived mesDA neurons with healthy controls. **e)** Graphical representation of lipid class concentration. Significance was calculated with Unpaired T-test analysis. p-value was set to statistically significant at <0.05. * Corresponds to <0.05, **<0.01, ***<0.001

Interestingly, apart from GM3 and CAR which were significantly down– and up-regulated, respectively, in the most of PD cell lines, we observed a highly heterogeneous expression of the lipid classes in the PD group, suggesting that different mutations could impact on distinct aspects of lipid metabolism (**Supplementary Fig. 5**).

In particular, the most dysregulated patient-derived neurons were from PD1, PD4 and PD6, carrying mutations in *KIF21B*, *SLC6A3*, *HMOX2, TMEM175* and *AIMP2* in which increased levels of FA, PC, PC-O, PE, Cer and DG were observed (**Supplementary Fig. 5**). Glycerosphingolipids (HexCer) were mainly altered in patients PD5 and PD6 with mutations in *LRRK2*, *TOMM22* and *RHOT2* (PD5) and *AIMP2* and *TMEM175* (PD6) (**Supplementary Fig. 5**). PD2 and PD3 cells showed a minor dysregulated lipid profile, which was associated with electrophysiological properties similar to HS (**Figs. 3, 4, Supplementary Fig. 5**).

We also analysed, across the different cell lines, the lipid species that contributed most to group separation in the PLS-DA model, as identified by the Variable Importance in Projection (VIP) analysis (**Fig. 5d**). Notably, several sulfoglycosphingolipids (SHexCer 41:7;O2, SHexCer 42:8;O2, SHexCer 39:7;O3) showed consistent differences in PD1, PD4, PD5, and PD6, whereas Triglyceride (TG) species showed prominent differences mainly in PD1, PD4, and PD6 (**Supplementary Fig. 6**). However, PD1, PD2 and PD3 also reflected reduced levels of LPC 20:1, while PD1 was the only one showing increased levels of DG 47:12 (**Supplementary Fig. 6**).

Univariate differential abundance analysis identified a defined set of lipids that were significantly modulated across the PD cell lines compared with. In particular, several neutral lipids were consistently altered, with a predominance of up-regulated TG and DG species, including DG 40:6 (FC = 48.6, p = 0.008), DG 41:0 (FC = 46.9, p = 0.0003), and TG 50:5|TG 16:1_16:2_18:2 (FC = 44.2, p = 0.007), indicating a robust remodelling of neutral lipid storage/turnover in PD cells. Beyond neutral lipids, several membrane lipids were also increased, including Phosphatidylglycerol (PG) 36:3|PG 16:0_20:3 (FC = 31.8, p = 0.005) and ether phospholipids such as PC O-40:6 (FC = 15.5, p = 0.0002), as well as selected sphingolipids (Cer 41:2;O2|Cer 18:1;O2/23:1, FC = 9.7, p = 0.006; SHexCer 39:1;O3, FC = 3.9, p = 0.002). Conversely, the down-regulated profile was dominated by sphingolipids and anionic phospholipids, including SHexCer 39:8;O3 (FC = 0.2, p = 1.5E-05), SHexCer 39:7;O3 (FC = 0.15, p = 6.3E-05), GM3 34:1;O2 (FC = 0.24, p = 5.2E-05), multiple Sphingomyelin (SM) species (e.g., SM 42:3;O2, FC = 0.48, p = 0.001), and phospholipids such as Phosphatidylinositol (PI) 36:4|PI 16:0_20:4 (FC = 0.52, p = 0.001) and Phosphatidylserine (PS) 32:1|PS 16:0_16:1 (FC = 0.29, p = 0.0004), with additional marked decreases in cardiolipin-related species (dilysocardiolipin (DLCL) 38:4, FC = 0.16, p = 0.0002) and ether PE/PC (e.g., PE O-34:5|PE O-14:1_20:4, FC = 0.038, p = 0.003; PC O-34:4|PC O-18:4_16:0, FC = 0.033, p = 0.0007).

Collectively, these results demonstrate true modulation of specific lipid pathways supported by statistical significance, rather than solely multivariate discrimination.

### Proteins dysregulated in patient-derived dopaminergic neurons were expressed in Sox6/AGTR1 positive nuclei and were altered in PD patients

To perform a comprehensive analysis of the presence of altered metabolic pathways in PD patients we investigated the proteomic profile in the same pool of iPSC-derived dopaminergic neurons in which we conducted the lipidomic analysis. Proteomic analysis identified 7844 proteins, quantified across all the samples. Statistical analysis showed the presence of hundreds of modulated proteins when comparing data from PD patients versus healthy subjects.

We identified 394 down-regulated proteins (FC ≤ 0.70; p ≤ 0.01) and 535 significantly upregulated proteins (FC ≥ 1.3; p ≤ 0.01) by comparing data from PD versus controls cell lines (**Fig. 6a).**

**Figure 6:**
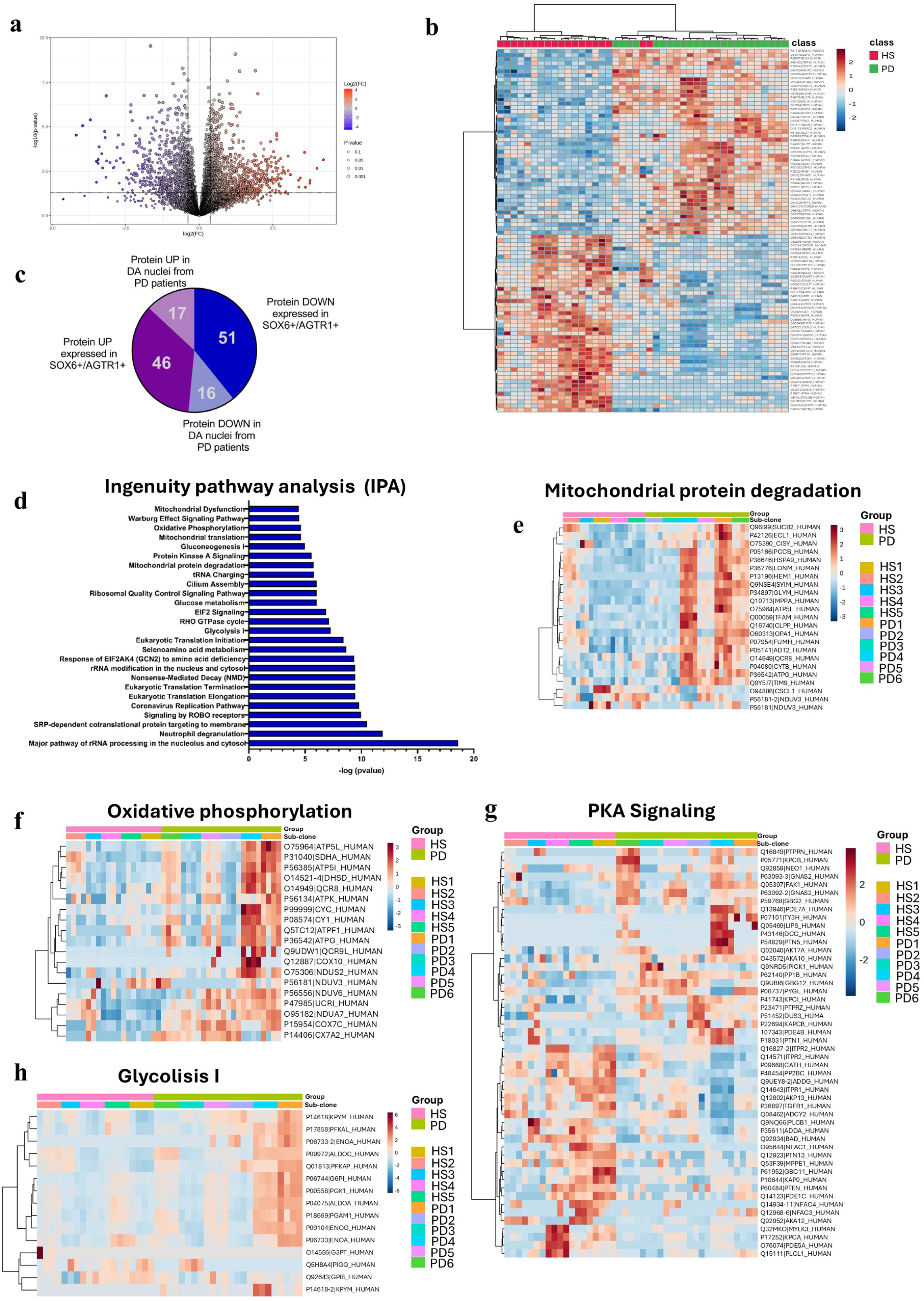
Proteomic analysis identified a high number of deregulated proteins (DEPs) in PD-derived mesDA neurons compared to the healthy subjects. **a)** Volcano plot showed the distribution of the most significant altered proteins comparing PD-derived mesDA neurons vs HS. Proteins showing statistically significant differences in expression are located in the top right (upregulated) and top left (downregulated) quadrants. The black line represents the p-value threshold set to p < 0.01. **b)** Heat map representation of the most 50 differentially expressed proteins highlighting the clusters of the two groups of analysis when comparing PD-derived mesDA to HS**. c)** Graphical representation of DEPs detected in PD-derived mesDA, up– (in red) and down– (in blue) regulated proteins that were also deregulated in SOX6+/AGTR1+ and in DA nuclei. Data from SOX6+/AGTR1+ and DA nuclei were available from the human single-cell RNA sequencing available dataset (https://singlecell.broadinstitute.org/single_cell/app/genes). **d)** Graphical representation of the most 20 enriched pathways from IPA analysis. **e-g**) Heat map representation of the most dysregulated pathways within the PD-derived mesDA when compared to the HS.

To focus on proteins related to neuronal physiology and potentially related to PD, we conducted a comprehensive literature search, by surveying the PubMed tool and we identified 62 out of the 394 down-regulated proteins expressed in neurons and related to PD (**Table S4**). However, considering the specific loss of the dopaminergic neurons of the Substantia Nigra pars compacta in PD patients, we tested the expression of the genes encoding for the 62 down-regulated proteins in the SNpc cells, by analyzing the human single-cell RNA sequencing available dataset [68] (https://singlecell.broadinstitute.org/single_cell/app/genes). We identified 51 out of 62 PD related genes expressed in Sox6/AGTR1 positive nuclei, which are those more sensible to neurodegeneration [68] **(Table S4; Fig. 6c)**. 16 proteins (AMPN (gene: *ANPEP*), AT1A2 (*ATP1A2*), GBG11 (*GNG11*), IP3KB (*ITPKB*), NUCKS (*NUCKS1*), UACA (*UACA*), CATH (*CTSH*), CATL1 (*CTSL*), CERS2 (*CERS2*), DGKQ (*DGKQ*), GIPC1 (*GIPC1*), GSDMD (*GSDMD*), PAG15 (*PLA2G15*), TRADD (*TRADD*), IFIT2 (*IFIT2*), TPPP3 (*TPPP3*)) were also down-regulated in DA nuclei from PD patients compared to controls and 6 of them (AMPN, AT1A2, GBG11, IP3KB, NUCKS, UACA) were also reduced in DA nuclei from Lewy body disease (LBD) patients (https://singlecell.broadinstitute.org/single_cell/app/genes) (**Table S4).**

Subsequently, we analyzed the expression of the 62 proteins in each of the six PD patients, individually. Interestingly, we disclosed 10 proteins that were significantly down-regulated in at least three PD patients, but only the Calpastatin protein ICAL (*CAST*) was significantly downregulated in all PD cells (**Tables S4**).

Among the most up regulated proteins we identified 49 proteins expressed in neuronal cells and related to PD (by PubMed survey), and 45 of them were expressed in Sox6/AGTR1 positive nuclei (**Table S5; Fig. 6c**). Interestingly, 17 up-regulated proteins (RASN (*NRAS*), CXCR4 (*CXCR4*), KAPCB (*PRKACB*), PP1B (*PPP1CB*), GRP75 (*HSPA9*), RALB (*RALB*), TFAM (*TFAM*), BCAT1 (*BCAT1*), SORL (*SORL1*), SESN2 (*SESN2*), TRAP1 (*TRAP1*), ILDR2 (*ILDR2*), GRK5 (*GRK5*), OPA1 (*OPA1*), C1QL3 (*C1QL3*), DDC (*DDC*), ELAV4 (*ELAVL4*)) were also increased in DA nuclei from PD patients compared to controls and 3 of them (RASN, CXCR4, KAPCB) were also increased in DA nuclei from Lewy body disease (LBD) patients (https://singlecell.broadinstitute.org/single_cell/app/genes) (**Table S5**).

Focusing on the data of the single PD patients we disclosed 32 proteins that were significantly up regulated in at least three PD patients, but only the protein LSM7 (*LSM7*) was significantly up-regulated in all PD cells (**Table S5**).

Altogether our data highlighted that the patients PD1, PD4 and PD6 showed a similar expression profile compared to PD2, PD3 and PD5, as also reported in lipidomic analysis and in electrophysiology studies (**Figs. 3-5**).

### Patient-derived dopaminergic neurons disclosed alteration of pathways involved in energy metabolism and mitochondrial processes

To identify altered pathways associated with the most dysregulated proteins identified in the proteomic analysis, we performed Ingenuity Pathway Analysis (IPA) by comparing protein levels in PD-derived neurons with those in control cells. IPA analysis identified about 199 significantly, dysregulated pathways (p < 0.01). Interestingly among the top 20 enriched pathways **(Fig. 6d)** we observed several processes related to cellular and mitochondrial metabolism and energy production, including: Warburg Effect Signaling Pathway, Oxidative Phosphorylation and Mitochondrial protein degradation **(Fig. 6d-f)**. Of interest is also the presence of deregulated proteins involved in Glucose metabolism, Gluconeogenesis and Glycolysis (**Fig. 6d,h**). However, we found that PD-derived cells, analyzed individually, exhibited a heterogeneous proteomic signature comprising molecules involved in the previously mentioned pathways. Indeed, PD1, PD4 and PD6 reflected a strong dysregulation of proteins such as UQCR10, COX7C and CAT involved into Oxidative Phosphorylation and ENO2, ALDOA/C, PKM, PFKL and PFKM which are key enzymes in regulating Glucose metabolism (**Fig. 6f,h Supplementary Fig. 7; Table S6**).

**Figure 7:**
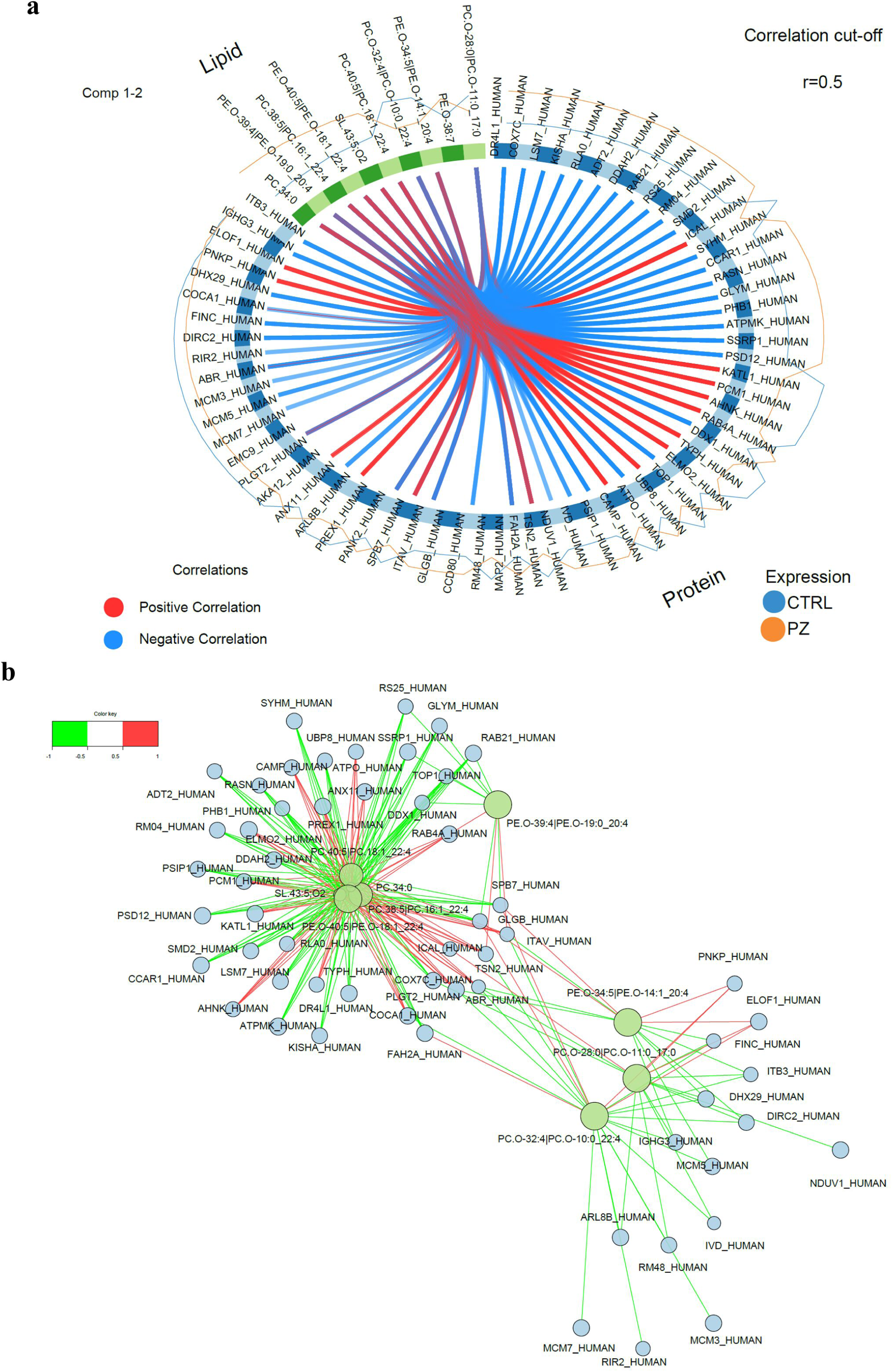
Integrative correlation analysis of proteome and lipidome data from PD-derived mesDA. **a)** Circos plot representing the averaged expression of proteins and lipids in each cell type (PD vs HS) and correlations between the features selected in each of the data blocks (protein features coloured in blue, and lipids features in green). **b)** Relevant network plot showing positive (red lines) and negative (green lines) correlations between features from each dataset.

Furthermore, when considering pathways closely related to mitochondrial stability, PD1, PD4 and PD6 displayed a marked alteration of MFN1, OPA1 and LONP1 belonging the mitochondrial protein degradation pathway (**Supplementary Fig. 7; Table S6**). Among the most enriched pathways, Protein Kinase A (PKA) signalling also showed significant enrichment of 48 dysregulated proteins in the PD group compared to controls **(Fig. 6d,g)**. In particular, PD1, PD4, and PD6 exhibited significant alterations in KPCB, TY3H, PTN5, AKAP13, and ADDA/G, confirming the presence of a distinct molecular signature potentially resulting from a patient-specific background **(Table S6)**. Conversely, PD2, PD3, and PD5 exhibited a completely different proteomic signature. Specifically, PD2 showed enrichment of pathways related to cell cycle regulation and DNA synthesis. PD3 displayed dysregulation of proteins involved in mRNA processing and cytoskeletal organization, whereas PD5 showed enrichment of altered proteins implicated in ferroptosis and branched-chain amino acid metabolism (**Supplementary Fig. 8**). These results evidenced the presence of an intricate proteomic signature within the PD group, suggesting a potential link with the different genetic background.

### Expression analysis of dysregulated proteins in mesDA neurons

To gain deeper insight into the proteomic dysregulation observed in the PD group, we investigated whether the alterations detected at the protein level were also reflected at the transcriptional one.

To this end, we analyzed by quantitative PCR, the expression of a selected panel of genes encoding the most dysregulated proteins impacting on the most enriched cellular pathways.

Considering that the expression of the selected proteins was highly heterogeneous among the PD patients when analyzed individually, each analysis was performed by comparing each PD patient with the HS group. Notably, although a high degree of heterogeneity in expression was observed among the different PD lines, we identified a highly concordant expression profile, at both the protein and transcriptional levels, for CAT, UQCR10, PKM, BCAT, TPPP3, and CXCR4 **(Supplementary Fig. 7)**. Conversely, the reduced levels of CAST protein detected in all PD patients, as well as the increased levels of COX7C, were not confirmed at the mRNA level **(Supplementary Fig. 7)**. Similarly, discrepancies between mRNA and protein levels were also observed for OPA1 and MFN1 **(Supplementary Fig. 7)**.

Overall, these findings suggest the involvement of differential transcriptional and post-transcriptional regulatory mechanisms across the various PD cell lines.

### Integrative correlation analysis of proteome and lipidome in iPSC-derived dopaminergic neurons

The correlations between proteomics and lipidomics data were evaluated using a supervised multi-omics integrative analysis which maximize the correlations between different types of omics and identify key molecules which can discriminate different sample groups. This integration was used to investigate the relationship between proteins and lipids across iPSC-derived dopaminergic neurons from PD patients and healthy subjects. The integrated model selected proteins and lipids that best explain PD-related variations in dopaminergic neurons. The general correlation between proteome and lipidome was high (0.63), suggesting common information among multi-omics data. In **Figure 7a,b** the circus and network plots depict correlations between modulated proteins and lipids are reported. A strong negative correlation between proteins and lipids was found, indicating a functional or physical lipid–protein interaction.

To explore the differences in the protein-lipid association, a topological analysis of protein-lipid correlation networks was conducted using all strongly correlating lipid-protein pairs based on significant strong Pearson correlations of r > |0.5|. A cluster of coregulated features strongly relevant to the latent components of the multi-omics dataset was identified, which might be a potential characteristic of dopaminergic neurons in PD patients (**Fig. 7b)**. The topological network is centred around two main lipid clusters: the first cluster included PC 40:5, PC 34:0, PC 38:5, PEO-40:5 and Saccharolipids (SL) 43:5 while the second one comprises PC O 32:4, PC O 28:0 and PE O 34:5.

Notably, several positive correlations emerge among the first cluster, including AHNAK, TYPH, COCA1, ABR, PLGT2, ICAL, RAB4A, KATL1, ELMO2, CAMP, PREX1, ANX11, PCM1, and ITAV. 26 proteins were negatively correlated: RAB21, GLYM, RS25, TOP1, SSRP1, DDX1, SYHM, ATPO, RASN, ADT2, RM04, DDAH2, PSIP1, PSD12, SMD2, CCAR1, FAH2A, KISHA, DR4L1, LSM7, RLA0, ATPMK, COX7C, SPB7 and GLGB.

Regarding the second cluster, only two proteins were positively correlated with lipids PNKP and ELOF1, while most of the proteins, namely FINC, ITB3, DHX29, DIRC 2, IGHG3, MCM5, MCM3, MCM7, NDUV1, ARL8B, IVD, RM48, and RIR2 were negatively correlated. Overall Network-based visualization of strongly correlating protein-lipid pairs highlighted two principal lipid hubs dominated by phosphatidylcholines and ether-linked glycerophospholipids, suggesting that these lipid classes are central to neuronal membrane structure. Moreover, this analysis also led to the identification of protein-lipid interactions that could not be detected by analysing each omics layer alone.

### Genetic association analysis revealed new PD risk genes

mesDA neurons derived from PD patients revealed distinct molecular signatures at the cellular level, characterized by dysregulation of proteins involved in energy and mitochondrial metabolism pathways as well as PKA signaling. To strengthen the disease relevance of these findings, we investigated whether genes encoding the most dysregulated proteins within these pathways were associated with PD.

We selected a panel of candidate genes, including *ADD1* and *AKAP13*, involved in PKA signaling; *LONP1*, *OPA1*, *CAT*, and *COX7C*, associated with mitochondrial function and oxidative stress pathways; and *ENO2*, *PKM*, *PFKL*, and *PFKP*, related to glycolysis and glucose metabolism pathways (**Tables S4-S6**). Furthermore, we included *BCAT1*, *CXCR4*, *LSM7*, *TPPP3*, and *CAST*, which encode the most up– and downregulated proteins identified in our analysis (**Tables S4-S6**).

To this end, genomic data were extracted from WES data from a large cohort of 833 PD patients and 688 controls. 500 variants with Minor Allele Frequency (MAF) > 0.01 (MAF was referred to our internal investigated cohort) were identified in the genomic regions, spanning from start to end (based on Genome Reference Consortium Human Build 38 (GRCh38)), of the selected genes.

Case-control association analysis was performed using PLINK2 software, applying logistic regression under an additive genetic model and adjusting for sex and the first 10 principal components (PCs) to account for genetic differences among individuals (**Supplementary Fig. 9a,b**). Interestingly, we identified 56 genetic variants located in intronic and exonic regions, as well as in regulatory genomic regions (3′-UTR and 5′-UTR), that were associated with PD (**Table S7**).

The analysis of the genes involved into PKA signaling pathway disclosed missense variants in *ADD1* and *AKAP13* associated with PD. The most associated variants included rs4961 (p.Gly460Trp; p = 0.0001; Odd Ratio (OR) = 0.67) and rs4963 (p.Ser648Cys; p = 0.0001; OR = 0.66) in *ADD1* and rs745191 (p.Gly624Val; p = 0.005; OR = 1.24), rs2241268 (p.Gly2457Ser; p = 0.0002; OR = 1.33) in *AKAP13* gene. One intronic variant rs3214982 (p = 0.0001; OR = 0.75) and one missense variant rs11085147 (p.Arg241Gln; p = 0.0003; OR = 0.54) were identified in the mitochondrial genes *OPA1* and *LONP1*, respectively. The most associated variants in genes related to glucose metabolism were: rs139729747 (p = 0.002; OR = 2.55) in the splicing regions of *ENO2*; rs3831402 (p = 3.2E-27; OR = 2.75) in the regulatory region in the first intron of *PFKL*; and rs375079219 (p = 3.4E-10; OR = 2.72) in the intronic region of *PFKP*. Interestingly, among the most dysregulated proteins we disclosed rs2680880 (p = 1.99E-11; OR = 0.57) in the 5’-UTR region of *CXCR4* gene and rs57890546 (p = 0.002; OR = 1.46) in the intronic region of *CAST* gene **(Table 2; Table S7)**.

**Table 2.**
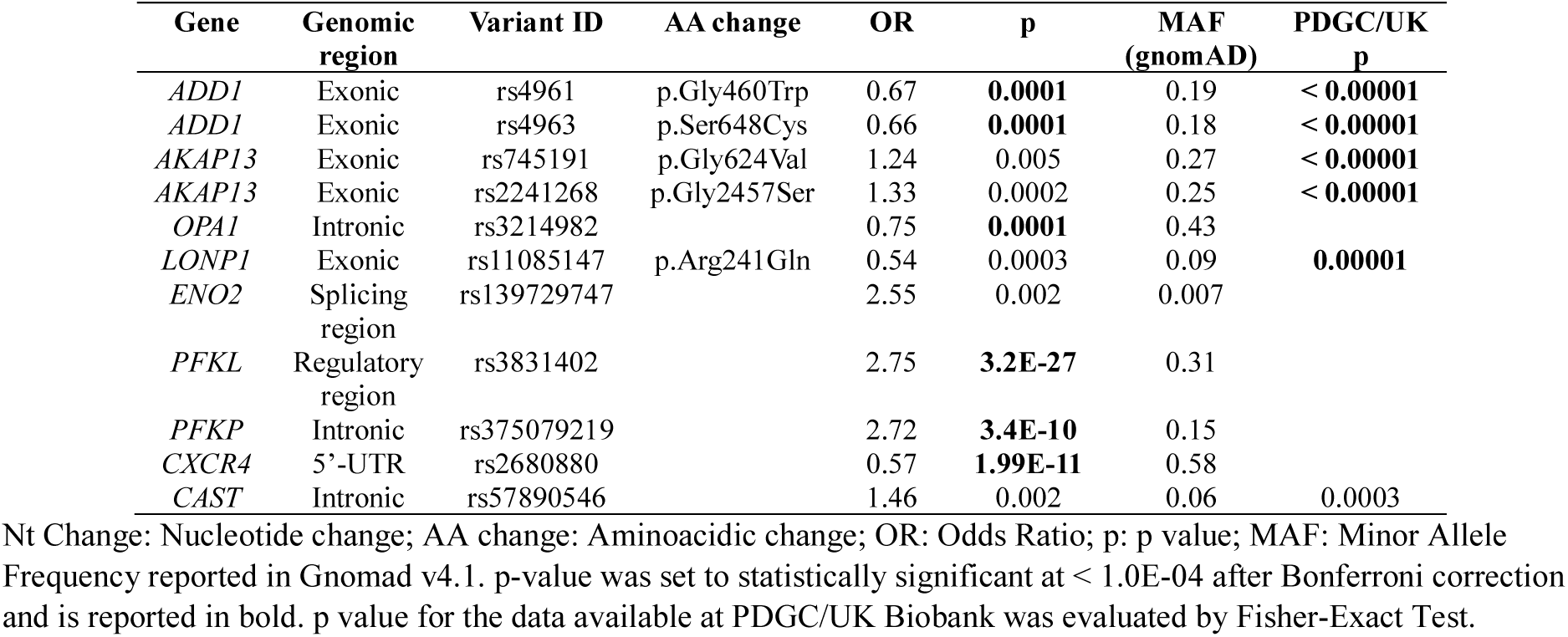
Genetic association analysis highlighted new variants in genes encoding for deregulated proteins associated with PD.

| Gene | Genomic region | Variant ID | AA change | OR | p | MAF (gnomAD) | PDGC/UK p |
| --- | --- | --- | --- | --- | --- | --- | --- |
| *ADD1* | Exonic | rs4961 | p.Gly460Trp | 0.67 | **0.0001** | 0.19 | < **0.00001** |
| *ADD1* | Exonic | rs4963 | p.Ser648Cys | 0.66 | **0.0001** | 0.18 | < **0.00001** |
| *AKAP13* | Exonic | rs745191 | p.Gly624Val | 1.24 | 0.005 | 0.27 | < **0.00001** |
| *AKAP13* | Exonic | rs2241268 | p.Gly2457Ser | 1.33 | 0.0002 | 0.25 | < **0.00001** |
| *OPA1* | Intronic | rs3214982 |  | 0.75 | **0.0001** | 0.43 |  |
| *LONP1* | Exonic | rs11085147 | p.Arg241Gln | 0.54 | 0.0003 | 0.09 | **0.00001** |
| *ENO2* | Splicing region | rs139729747 |  | 2.55 | 0.002 | 0.007 |  |
| *PFKL* | Regulatory region | rs3831402 |  | 2.75 | **3.2E-27** | 0.31 |  |
| *PFKP* | Intronic | rs375079219 |  | 2.72 | **3.4E-10** | 0.15 |  |
| *CXCR4* | 5'-UTR | rs2680880 |  | 0.57 | **1.99E-11** | 0.58 |  |
| *CAST* | Intronic | rs57890546 |  | 1.46 | 0.002 | 0.06 | 0.0003 |

Nt Change: Nucleotide change; AA change: Aminoacidic change; OR: Odds Ratio; p: p value; MAF: Minor Allele Frequency reported in Gnomad v4.1. p-value was set to statistically significant at < 1.0E-04 after Bonferroni correction and is reported in bold. p value for the data available at PDGC/UK Biobank was evaluated by Fisher-Exact Test.

Genetic variants resulting associated after Bonferroni correction for multiple testing (number of variants = 500; p < 1.0E-04) included rs4961, rs4963, rs3214982, rs3831402, rs375079219 and rs2680880 in *ADD1, LONP1, PFKL, PFKP* and *CXCR4* genes **(Table 2)**.

Furthermore, to better highlight the relevance of the associated variants, we performed a case-control association analysis in a larger independent cohort comprising 4,586 PD patients from the Parkinson’s Disease Genetic Consortium (PDGC) and 43,989 individuals from the general population enrolled in the UK Biobank, without adjusting for sex or genetic stratification. The analysis comparing the PDGC cohort with the UK Biobank was performed using Fisher’s exact test, as only aggregated genotype data were available for this study cohort (https://pdgenetics.shinyapps.io/VariantBrowser/). Among the most associated variants, 9 variants (rs4961 (p < 0.00001); rs4963 (p < 0.00001); rs745191 (p < 0.00001); rs2241268 (p < 0.00001); rs11085147 (p = 0.00001); rs57890546 (p = 0.0003) were found in the PDGC/UK dataset and Fisher-Exact test confirmed a strong association with PD (**Table 2**). Moreover, to determine whether these genetic variants were associated with differences in allelic expression in human brain tissues, we queried the GTEx portal, a public dataset containing Quantitative Trait Loci (QTL) data from human-derived brain tissues (https://gtexportal.org/home/). We found that rs4961, rs4963, rs745191, rs2241268, rs3214982, rs11085147, rs3831402, and rs57890546 were associated with differential allelic expression in brain tissues (**Tables S8–S15**). The most associated alleles were the protective allele rs11085147-T (p.241Gln) in the *LONP1* gene and the risk allele rs3831402-G in the *PFKL* gene, both of which were associated with a highly significant reduction of *LONP1* and *PFKL* transcript levels, respectively, across all brain tissues (**Supplementary Fig. 10 a and b; Table S13 and 14**).

These findings further support the validity of our approach in detecting novel risk variants associated with PD by using multi-omics and multilayers analyses.

## Discussion

In the present study, we investigated the functional consequences of the combined presence of rare variants in novel PD genes, an aspect that remains poorly characterized, particularly in disease-relevant cellular contexts.

We adopted an integrative approach, examining the rate of differentiation into post-mitotic dopaminergic neurons, electrophysiological properties, and lipidomic and proteomic profiles, in order to achieve a comprehensive molecular characterization of hiPSC-derived mesDA neurons carrying newly identified rare genetic variants associated with PD.

We generated and extensively characterized a panel of patient-derived hiPSC lines harbouring distinct combinations of PD-associated genetic variants and successfully differentiated them into ventral midbrain dopaminergic neurons. Notably, all lines retained robust pluripotency, genomic stability, and comparable efficiency in acquiring a dopaminergic identity, as reflected by the formation of FOXA2⁺/LMX1⁺ progenitors and mature TH⁺ neurons. The absence of significant differences in differentiation efficiency between PD– and control-derived lines suggests that, under the conditions used, these genetic variants do not substantially interfere with early lineage specification or neuronal maturation. This observation aligns with previous reports in iPSC-based PD models, where disease-related phenotypes typically manifest at later stages of neuronal maturation or under stress conditions, rather than during initial differentiation[69]. Importantly, the consistent generation of mature dopaminergic neurons across all lines establishes a robust and well-controlled platform for downstream functional investigations.

To achieve a comprehensive characterization of the generated mesDA neurons, we applied patch-clamp analysis and omics approaches, including lipidomic and proteomic profiling, in these cell lines. Interestingly, dopaminergic neurons derived from different PD patients exhibited distinct electrophysiological profiles. PD1 (*KIF21B*), PD4 (*SLC6A3, HMOX2*), PD5 (*LRRK2*, *RHOT2*, *TOMM22*) and PD6 (*AIMP2*, *TMEM175*) neurons showed profound alterations in excitability, while PD2 (*ZSCAN21*, *ANKK1*) and PD3 (*TMEM175*, *TVP23A*) neurons were less affected when compared to HS. Furthermore, PD4 carrying a pathogenic mutation in DAT (*SLC6A3*) also exhibited functional impairments in excitatory glutamatergic synaptic transmission, suggesting that this phenotype, involving an altered dopamine transporter, destabilizes the overall neuronal signalling more deeply. These data correlate well with lipidomic and proteomic measurements: both profiles are highly dysregulated for PD4, while those of PD2 and PD3 are more similar to HS. This observation indicates a relevant direct relationship between lipidomic, proteomic, and functional alterations in patient-derived hiPSC dopaminergic neurons. Among all genotypes, PD3 is the only one to exhibit increased neuronal excitability; this finding may indicate functional compensation and is associated with a less severe clinical phenotype.

Since lipid metabolism have been closely linked to PD pathophysiology [70–72], we aimed to explore whether PD-derived mesDA neurons could reflect the presence of a perturbed lipid composition, when compared to HS. Effectively, our results supported the presence of a distinct lipid profile in PD-derived mesDA neurons showing a wide increase in most of the lipid classes. Specifically, PD neurons showed increased levels of acylcarnitines and fatty acids. Perturbations in these lipids have been reported to affect energy production, mitochondrial homeostasis, fatty acid oxidation, and cellular stress, ultimately leading to toxic accumulation and impaired dopamine signaling. This imbalance disrupts energy production, causing mitochondrial enlargement and oxidative stress, impacting dopamine neuron health and function[73]. Moreover, increased levels of glycerophospholipids, including PC and PE, have previously been reported in the plasma and serum of PD patients[47,74,75]. In this study, we observed this upregulation at the cellular level, potentially linking it to disruptions in plasma and mitochondrial membrane stability, as also suggested by the altered electrophysiological properties of PD-derived mesDA neurons. Interestingly, PD3 which is the only PD cell line evidencing an increased neuronal excitability, reflected a completely different lipidomic profile compared with the other PD cell lines. Additionally, we disclosed an upregulation of Cer, HexCer and SHexCer, members of the Sphingolipid class, within the PD group. These lipids are reported not only as major components of the plasma membrane but also as key signaling molecules involved in several cellular processes[76,77]. Furthermore, they are involved in neuronal development and function[77], highlighting the significance of their alteration in neurodegenerative conditions.

Interestingly, Sulfoglycosphingolipids, or sulfatides, are a type of glycosphingolipid found in neurons that are critical for neural development, function, and integrity. They contribute to the complexity of the neuronal plasma membrane and play roles in cell signaling, adhesion, and the formation of specialized structures like synapses[78]. Dysregulation of sulfoglycosphingolipid metabolism has been linked to neurodegenerative diseases[79–81].

Conversely, despite the overall increase in most lipid classes, we observed a marked decrease in Ganglioside 3 levels in PD patients. GM3 is well known to be involved in the maintenance of cell membrane stability, which is strongly affected in PD, and its reduced expression has already been reported in PD[82–84]. Furthermore, to determine whether the observed lipidomic dysregulation was present in all PD-derived mesDA neurons or specific to certain lines, we analyzed the lipid levels in each PD-derived cell line, revealing variability in their abundance within the group. Interestingly PD1, PD4 and PD6, carrying mutations in *KIF21B*, *SLC6A3*, *HMOX2*, *TMEM175* and *AIMP2*, exhibited alterations in most lipid classes compared with HS, potentially indicating a mutation-specific phenotype affecting lipid metabolism. Conversely, acylcarnitines were the only lipid class altered across all PD neurons, including PD2, PD3, and PD5, consistent with evidence that these lipids play a critical role in brain function and PD pathophysiology [85].

Consistent with the distinct molecular, electrophysiological, and lipidomic profiles observed across PD-derived mesDA neurons, proteomic analysis revealed varying patterns of protein deregulation. We first demonstrated that proteins expressed by SOX6/AGTR1⁺ neurons and associated with PD are strongly dysregulated in PD-derived cell lines, confirming the validity of our cellular model. Considering the entire group of PD-derived cell lines, our results showed that Calpastatin (CAST) protein is markedly downregulated in all analyzed PD patient–derived cell lines. Calpastatin is an endogenous inhibitor of calpains, which are intracellular, Ca²⁺-dependent cysteine proteases ubiquitously expressed in most tissues and organs, including the brain, and exhibit tightly regulated protease activity[86]. Among their target, Calpains have been described as regulator of alpha-synuclein degradation and, consequently, their changes in terms of activity could be implicated into neurodegeneration[87]. Moreover, in accordance with our data, it has been reported downregulation of Calpastatin and increased levels of Calpains in the mesencephalon of PD patients[88], highlighting its potential relevance as new target in relation to PD pathophysiology. On the other hand, the entire group of PD-derived neurons also showed significant increase in LSM7, CXCR4 and BCAT1 proteins, suggesting their involvement in PD pathophysiology. These findings are in accordance with previous studies demonstrating the presence of genetic risk variants in LSM7 [89], as well as several evidence described CXCR4 as relevant link between neuroinflammation and PD [90]. Additionally, the alteration of BCAT1, which is an enzyme involved in the metabolism of Branched chain amino acids, is in accordance with previous evidence reporting perturbations of these amino acids in the plasma of PD patients carrying specific genes mutations [47,75].

Furthermore, to gain insight into the most enriched pathways in our cellular models, IPA analysis detected several pathways that were differentially perturbed by deregulated proteins across the PD-derived cell lines.

The PKA signaling pathway showed clear clustering between PD and HS groups, suggesting that proteins involved in this process are differentially expressed. PKA plays a central role in the regulation of dopaminergic neuron function, survival, and plasticity, primarily acting downstream of cyclic AMP (cAMP) generated by dopamine receptor activation. In the nigrostriatal system, PKA integrates signals from dopamine D1/D2 receptors, neurotrophic factors, and neuromodulators, thereby influencing transcriptional programs, mitochondrial activity, and synaptic function [91]. Dysregulation of this pathway has increasingly been implicated in the pathogenesis of PD [92–94]. Our results are in line with previous studies using human iPSC-derived dopaminergic neurons highlighting PKA signaling as a dysregulated pathway in patient-derived models. This is particularly relevant in the context of genetically heterogeneous PD, where distinct rare variants may perturb PKA signaling through different upstream mechanisms while converging on shared downstream phenotypes.

On the other hand, we also reported that PD-derived neurons exhibited deregulated levels of proteins involved in the most relevant pathways related to energy production. In this context, we demonstrated that PD1, PD4 and PD6 were the most affected cell lines showing alterations in key enzymes involved in glycolysis (ENO2, ALDOA/C, PKM, PFKL and PFKM), oxidative phosphorylation (CAT, UQCR10, COX7C) and mitochondrial stability (OPA1, MFN1). Interestingly, most of these enzymes were deregulated at the protein level but not at the transcriptional level, potentially reflecting impaired post-transcriptional regulation in PD patients carrying mutations in *KIF21B, SLC6A3, HMOX2, TMEM175* and *AIMP2*.

Our previous findings were further supported by supervised multi-omics integration analysis, which extracts latent components that maximize shared and correlated information across omics layers while selecting discriminant features that best separate phenotypic groups. This approach enabled the identification of coupled protein-lipid signatures that are not apparent when each omics layer is analyzed individually. The relatively high global correlation (r = 0.63) between the selected proteomic and lipidomic components supports the presence of substantial common biological variation captured across platforms and is consistent with coordinated remodelling of cellular programs that affect both protein abundance and membrane composition. Network-based visualization of strongly correlating protein-lipid pairs highlighted two principal lipid hubs dominated by phosphatidylcholines and ether-linked glycerophospholipids, suggesting that these lipid classes are central to neuronal membrane structure [95,96]. Notably, a recent ultrastructural work has shown that Lewy pathology is enriched in membranous material, strengthening the rationale for interrogating protein-lipid co-variation in disease-relevant neurons [97]. Biologically, the strong negative correlations observed between subsets of proteins and lipids may reflect compensatory responses to membrane remodelling, shifts in lipid availability, or changes in trafficking and turnover of membrane-associated protein complexes. The co-variation of a phosphatidylcholine/plasmalogen-rich cluster with proteins involved in cytoskeletal organization, adhesion, and Rab-mediated membrane trafficking is consistent with the bidirectional links between vesicle trafficking and lipid homeostasis emphasized in synucleinopathies [70,98]. Conversely, the second ether-lipid cluster showing predominantly negative associations with proteins involved in nucleic acid metabolism and mitochondrial functions could indicate broader stress responses or altered bioenergetic demands in PD neurons, although these hypotheses require experimental validation.

Finally, our findings clearly demonstrate the presence of a heterogeneous electrophysiological and molecular phenotype in PD-derived mesDA neurons carrying different combinations of rare genetic variants. Importantly, while our results confirm alterations in key pathways closely related to PD and support the validity of our cellular model, they also reveal novel molecular markers, including Calpastatin, and several lipid alterations that have not previously been described at the cellular level. Based on this evidence we further investigate whether pathways/genes dysregulated in PD-derived neurons could be considered as genetic modifiers for PD. interestingly, we found missense and regulative variants significantly associated with PD in several genes, including *ADD1,* encoding a protein responsible for membrane cytoskeletal organization [99], *LONP1* encoding for a mitochondrial matrix protease [100]*, CXCR4* encoding for a chemokine receptor protein with broad regulatory functions in the immune system and neurodevelopment [101] and in two isoforms of Phosphofructokinase also referred as *PFKL* and *PFKM* which catalyse the first committed step in glycolysis [102]. Notably, some of these genes have already been described associated to neurodegenerative disorders. Indeed, *ADD1* has been reported as new risk locus to PD in larger case-control studies [103], while CXCR4 has been showed to enhance the microglial and astroglial activation of MPTP-lesioned mice [104]. LONP1 is a key player involved in the degradation of matrix mitochondrial proteins including PINK1. Mutations in *LONP1* gene are responsible for the CODAS syndrome with Cerebral, Ocular, Dental, Auricular and Skeletal abnormalities[105]. Recently a bi-allelic mutation (c.2282 C > T, (p.Pro761Leu) in *LONP1* has been associated to neurodegeneration with deep hypotonia and muscle weakness, severe intellectual disability and progressive cerebellar atrophy [106]. These observations clearly indicate a relationship of LONP1 with central nervous system abnormalities and neurodegeneration.

These results support the validity of our approach in capturing disease-relevant pathways and in identifying novel modifiers PD genes. We also found that the most strongly associated alleles were the protective rs11085147-T (p.241Gln) in the *LONP1* gene and the risk rs3831402-G in the *PFKL* gene, both of which were associated with a highly significant reduction in expression in brain tissues. These findings strongly support our hypothesis that, among the genes analyzed, *LONP1* and *PFKL* could represent novel PD risk genes.

Altogether, our results support a model in which combinations of rare genetic variants contribute to PD pathogenesis by perturbing key cellular processes and affecting multiple proteins that are actively expressed and deregulated in dopaminergic neurons. Moreover, the dysregulation of lipids and proteins involved in energy metabolism pathways in PD neuronal cells is consistent with previous studies reporting alterations in circulating molecules associated with these processes in the serum of PD patients carrying rare genetic variants. In particular, we previously observed an enrichment of dysregulated metabolites involved in nicotinate–nicotinamide metabolism, short-chain fatty acid metabolism, and amino sugar metabolism in both male and female patients compared with healthy subjects [107].

Notably, our results support the hypothesis that the alterations detected at cellular levels could mirror the metabolic profile at circulating level and could improve the identification of novel risk factors/modifiers genes.

Furthermore, our data demonstrated that the extreme genetic heterogeneity observed in PD patients reflects the synergistic effect of multiple rare variants in different PD genes. For example, PD3 and PD6, each carrying a *TMEM175* mutation in combination with different variants in *TVP23A* (PD3) and *AIMP2* (PD6), exhibited completely divergent molecular phenotypes.

Finally, we demonstrated that multi-omics profiling in patient-derived hiPSC models may facilitate the identification of biomarkers and therapeutic targets in relation to specific genetic backgrounds.

## Conclusions

In conclusion, our study demonstrates that hiPSC-derived dopaminergic neurons from PD patients carrying rare genetic variants constitute a robust and informative model to investigate both shared and variant-specific mechanisms of disease. By applying a multi-omics approach, we provide new insights into the molecular heterogeneity of PD and underscore the value of patient-specific cellular models for advancing our understanding of complex neurodegenerative disorders and potentially expanding the application of future precision medicine.

## Declarations

### Ethics approval and consent to participate

All procedures involving human participants were approved by the Institutional Review Board of the IRCCS Neuromed Italy. The study protocols N°9/2015, N°19/2020, N°4/2023 have been registered in clinicaltrial.gov with the numbers NCT02403765 (Release Date: 04/01/2015), NCT04620980 (Release Date: 11/03/2020), NCT05721911 (Release Date: 30/01/2023).

Clinical investigations were conducted according to the principles expressed in the Declaration of Helsinki. Written informed consent was obtained from all participants.

The research was carried out following the recommendations set out in the Global Code of Conduct for Research in Resource-Poor Settings.

## Consent for publication

Not applicable

## Data availability

The mass spectrometry proteomics data have been deposited to the ProteomeXchange Consortium via the PRIDE partner repository with the dataset identifier PXD074629. Lipidomic dataset is reported in Table S3

## Author contributions

FC: Investigation, data analysis, writing, review & editing; GF: Investigation, data analysis; AC: Investigation, data analysis; MG: Investigation, data analysis; NGO: Investigation, data analysis; VFB: Investigation, data analysis; SK: data analysis; ADL: Investigation; MM: Investigation; GG: Investigation; FDA: Investigation; MRI: Investigation; CDA: Investigation; AFiorenzano: review & editing; TN: Data analysis; DL: Data analysis; SP: review & editing; NM: review & editing; KM: Investigation, data analysis, review & editing; SF: Data analysis, review & editing; MM: Data analysis, review & editing; AFico: Funding acquisition, supervision, investigation, data analysis, writing – review & editing. TE: Conceptualization, funding acquisition, project administration, supervision, data analysis, writing – review & editing.

## List of abbreviations

PD: Parkinson’s disease
hiPSC: human induced pluripotent stem cell
DA: dopaminergic
SNpc: Substantia Nigra pars compacta
HS: healthy subjects
AAO: age at symptom onset
MDS-UPDRS: Movement Disorder Society revised version of the Unified Parkinson’s Disease Rating Scale Part III
HY: Hoehn e Yahr
MoCA: Montreal Cognitive Assessment
WES: Whole exome sequencing
MNI: Mediterranean Neurological Institute
EUR-nfe cohort: healthy non-Finnish European population
PDGC: Parkinson Disease Genetic Consortium
UK: United Kingdom
PBMCs: peripheral blood mononuclear cells
PBS: phosphate-buffered saline
EDTA: Ethylenediaminetetraacetic acid
qPCR: quantitative PCR
ACTB: beta actin gene
ddPCR: Digital droplets
PCR IF: immunofluorescence
RT: room temperature
PCA: Principal Component Analysis
CNT: controls
MAF: Minor allele frequency
GTEx: Genotype-Tissue Expression
DAT: Dopamine Transporter 1
PSC: pluripotent stem cell
VM: ventral midbrain
GSK3i: glycogen synthase kinase 3 inhibitor
SHH-C24II: Sonic Hedgehog
mesDA: mesencephalic dopaminergic neurons
TH: Tyrosine Hydroxylase
Aps: action potentials
EPSCS: excitatory post-synaptic currents
PLS-DA: Partial Least Squares Discriminant Analysis
FC: Fold Change
CAR: Acylcarnitine
FA: Fatty Acids
Cer: Ceramide
HexCer: Hexosylceramide
SHexCer: Sulfoglycosphingolipid
PC: Phosphatidylcholine
PC-O: Ether-linked Phosphatidylcholine
PE: Phosphatidylethanolamine
DG: Diacylglycerols
TG: Triglyceride
PG: Phosphatidylglycerol
SM: Sphingomyelin
PI: Phosphatidylinositol
PS: Phosphatidylserine
DLCL: dilysocardiolipin
GM3: ganglioside GM3
VIP: Variable Importance in Projection
LBD: Lewy body disease
IPA: Ingenuity Pathway Analysis
GRCh38: Genome Reference Consortium Human Build 38
QTL: Quantitative Trait Loci

## Acknowledgments

The authors are grateful to all the patients, their caregivers, the Clinical Parkinson’s Disease Center of IRCCS Pozzilli and the PD biobank of IRCCS Neuromed and IGB-CNR for the kind cooperation with this study.

